# Dynamic Spatiotemporal Organization of Exocytosis During Cellular Shape Change

**DOI:** 10.1101/185249

**Authors:** Fabio Urbina, Shawn Gomez, Stephanie L. Gupton

## Abstract

Morphological elongation of developing neurons requires plasmalemma expansion, hypothesized to occur primarily via exocytosis. We posited that exocytosis in developing neurons and non-neuronal cells exhibit distinct spatiotemporal organization. We exploited TIRF microscopy to image VAMP-pHluorin mediated exocytosis in murine embryonic cortical neurons and interphase melanoma cells, and developed computer-vision software and statistical tools to uncover spatiotemporal aspects of exocytosis. Vesicle fusion behavior differed between vesicle types, cell types, developmental stage, and extracellular environment. Experiment-based mathematical calculations indicated that VAMP2-mediated vesicle fusion supplied excess material for the plasma membrane expansion that occurred early in neuronal morphogenesis, which was balanced in part by clathrin-mediated endocytosis. Spatial statistics uncovered distinct spatiotemporal regulation of exocytosis in the soma and neurites of developing neurons that was modulated by developmental stage, exposure to the guidance cue netrin-1, and the brain-enriched ubiquitin ligase TRIM9. In melanoma cells, exocytosis occurred less frequently with distinct spatial clustering patterns.

## Introduction

Exocytosis is a fundamental cellular behavior ubiquitous across eukaryotes and cell types. Vesicle fusion promotes secretion of biomolecules and insertion of transmembrane proteins and lipids into the plasma membrane, which can affect physiological processes including polarized growth and motility (Mostov et al., 2000; Winkle et al., 2014). Where and when exocytic vesicle fusion occurs therefore represents a critical point of regulation in cellular physiology. The minimal protein machinery required for fusion is the evolutionarily conserved SNARE complex (Söllner et al., 1993), which comprises a stable, tightly-associated bundle of four α-helical coiled-coils provided by three proteins. During exocytosis, one α-helix is provided by a vesicle (v)-SNARE, such as vesicle-associated membrane protein 2 (VAMP2, synaptobrevin), VAMP3, or VAMP7 (tetanus-insensitive VAMP) in mammals or snc1/2 in yeast (Galli et al., 1998; McMahon et al., 1993; Protopopov et al., 1993). Other α-helixes are provided by plasma membrane target (t)-SNAREs: syntaxin-1 and synaptosomal-associated protein 25 (SNAP25) in mammals or Sso1p/Sso2p and sec9 in yeast (Aalto et al., 1993; Brennwald et al., 1994; Söllner et al., 1993).

VAMP2, SNAP25 and syntaxin-1 were identified in brain, where they perform a critical role during synaptic vesicle fusion and neurotransmitter release. VAMP7 functions in SNARE-mediated exocytosis in both neurons and non-neuronal cells (Galli et al., 1998; Voets et al., 2016). Subsequent to synaptic vesicle release, rapid clathrin-dependent endocytic retrieval of membrane material and v-SNAREs maintains membrane homeostasis (Heuser and Reese 1973; Pearse, 1976). Perhaps less appreciated than synaptic exocytosis is the developmental exocytosis that occurs prior to synaptogenesis. The acquisition of the elongated morphology of developing neurons entails significant plasma membrane expansion, estimated at a 20% per day (Pfenninger, 2009). This is remarkable when compared to concomitant neuronal volume increases estimated at less than 1%. We previously demonstrated that constitutive SNARE-mediated exocytosis is required during neuritogenesis and axon branching (Gupton and Gertler, 2010; Winkle et al., 2014). We hypothesize this provides membrane material to the expanding plasma membrane, which can only stretch 2-3% before rupturing (Bloom et al., 1991), however whether SNARE-mediated exocytosis supplies sufficient material for additional membrane expansion has not been addressed. Studies also suggest that asymmetric exocytosis is linked to attractive axonal turning responses (Ros et al., 2015; Tojima et al., 2014, 2007) that are critical for axon guidance. As a number of neurological disorders are accompanied by disrupted neuronal morphology (Engle, 2010; Grant et al., 2012; Paul et al., 2007), the regulated exocytosis involved in appropriate neuronal morphogenesis is likely central to the formation and maintenance of a functional nervous system. However how exocytosis is spatially and temporally organized in developing neurons is not known.

To visualize exocytic vesicle fusion, here we exploited the pH-sensitive variant of GFP (pHluorin) attached to the lumenal side of a v-SNARE, which illuminates the occurrence of fusion pore opening between the acidic vesicular lumen and the neutral extracellular environment (Miesenböck et al., 1998). To this point, analysis of VAMP-pHluorin images has remained non-automated and time-intensive, delaying our understanding of this fundamental cellular behavior. Here we developed computer-vision software and statistical methods for unbiased detection and analysis of VAMP-pHluorin mediated exocytosis. This uncovered spatial and temporal organization and regulation of exocytosis in developing neurons that was distinct in soma and neurites, which was modulated by the developmental stage of the neuron as well as exposure to the axon guidance cue netrin-1. Mathematical estimations based on empirical findings suggest VAMP2-mediated exocytosis and clathrin-mediated endocytosis approximately describe the membrane expansion of developing neurons. Compared to neurons, interphase melanoma cells exhibited slower frequencies and a distinct organization of exocytosis.

## Results

### Automated identification and analysis of exocytosis

Whether exocytosis is sufficient for neuronal plasmallemal expansion, how fusion is organized spatially and temporally, and the mechanisms that regulate developmental exocytosis are not established. We imaged VAMP2-pHluorin or VAMP7-pHluorin in murine embryonic cortical neurons by widefield epifluorescence microscopy to visualize exocytic events on ventral and basal plasma membranes and by Total Internal Reflection Fluorescence (TIRF) microscopy to visualize exocytic events specifically on the basal plasma membrane (Figure S1A). The frequency of VAMP2-mediated exocytosis measured from TIRF images, reported per basal plasma membrane area, was ~1.4 times higher than measured in widefield (reported per 2x basal plasma membrane area, Figure S1B), whereas the frequency of VAMP7-mediated exocytosis was comparable between the two imaging modes (Figure S1B). VAMP2-pHluorin exhibited a higher background fluorescence on the plasma membrane than VAMP7-pHluorin (Figure S1C), which likely occluded identification of VAMP2-mediated events on the ventral plasma membrane. This was supported by mapping the density of exocytic events in cells imaged by TIRF and subsequently by widefield microscopy at two distinct focal planes (Figure S1D, black arrows). Since both basal and ventral plasma membranes are not within a focal plane, events on both membranes cannot be captured at sufficient temporal resolution, and the rate of estimated fusion in widefield microscopy was likely an underestimation. We conclude that the frequency of exocytosis is comparable between ventral and basal plasma membranes for both VAMP2 and VAMP7-mediated exocytosis (Figure S1B, D), which was captured by TIRF microscopy. For subsequent analysis, we relied on TIRF microscopy, which also offered increased signal-to-noise, and spatial resolution, reduced photobleaching and phototoxicity, and faster image acquisition.

To perform unbiased, efficient identification and analysis of exocytosis, we developed a computer vision-based software in MATLAB and R, outlined in Figure 1A, based on TIRF images of VAMP2-pHluorin. The algorithm we developed has advantages over previously reported semi-automated and automated exocytic detection algorithms (Díaz et al., 2010; Sebastian et al., 2006). This includes a more rigorous definition of an exocytic event, experimental validation using appropriate controls, application to cells of complex shapes, such as primary neurons (Figure 1A; Movie 1), and the inclusion of statistical analyses of multiple spatial and temporal parameters of fusion events across conditions, providing higher confidence conclusions. In the analysis pipeline, first the cell was identified and segmented from non-cellular background; a single cell mask was created, as cells did not significantly change shape or size over the course imaging (~2 min). Frames were convolved with a Gaussian filter followed by local application of an h-dome transform (Vincent, 1993), to detect bright, round-shaped objects inside the cell mask against local variations in fluorescence. Gaussian mixed models were fit to local fluorescent maxima in each frame (Bishop, 2006), allowing the detection of both diffraction-limited and larger fluorescent objects. To link the detected fluorescent objects that were part of the same exocytic event across frames, a Kalman filter was used (Kalman, 1960) (Figure 1B, S2A). We defined a small radius expected to encompass the expansion and/or movement centered on potential fusion events identified in the initial frame (t_0_). Using a cost matrix (Jaqaman et al., 2008), detected events that fell into the prediction radius were identified as the same object; if not, they were identified as a new object. The parameters of the cost matrix, including distance between predicted and detected locations, were determined experimentally (Figure S2B,C). The prediction radius was refined in subsequent frames (t_i+1_) based on prior behavior of detected objects. This iterative process continued through the sequence of images to build connected object tracks, which represented potential exocytic events. *Bona fide* exocytic events were defined as transient, non-motile (mean squared displacement <0.2 μm^2^) objects that reached a peak fluorescence intensity four deviations above the average local background intensity at t0, followed by a detectable exponential fluorescent decay (≥3 frames) with no limit on how long the events could last (Figure 1C, Bowser and Khakh, 2007). Tracks that did not meet these requirements were discarded.

**Figure 1.**
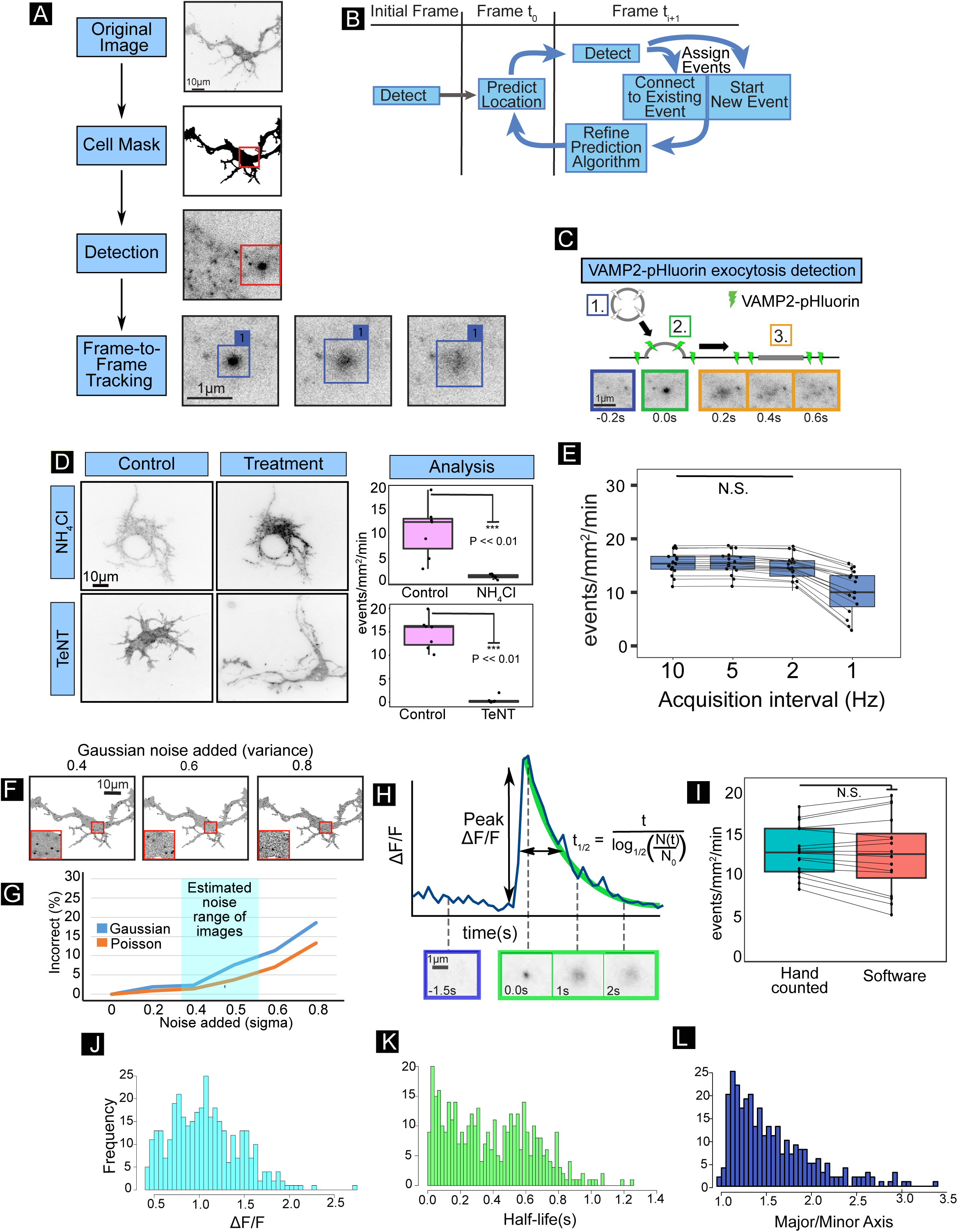
Automated identification, tracking and analysis of VAMP-pHluorin mediated exocytosis. **A)** Overview of automated software. A cell mask is first generated, followed by fluorescent bright spot maxima detection. A Kalman-filter based tracking matrix connects candidate events over time (Frame-to-Frame Tracking) to identify *bona fide* exocytic events. **B)** The tracking matrix was constructed using a Kalman filter to link a single exocytic event into a track over multiple frames. **C)** Schematic and TIRF image montage showing a typical identified exocytic event revealed by VAMP2-pHluorin fluorescence. Pre-fusion, VAMP2-pHluorin is not fluorescent (1, blue). Upon fusion pore opening (2, green), a bright fluorescent diffraction-limited spot appears. The VAMP2-pHluorin diffuses in the membrane over time (3, orange). **D)** Example minimum projections of TIRF time-lapse images of neurons before or after treatment with sNH_4_CI or Tetanus neurotoxin (TeNT). Graphs indicate the frequency of exocytosis detected by the automated image analysis software (n = 6 cells per condition). NH_4_Cl or TeNT treatment reduced the number of events detected (boxplot, median ± S.D.) See movie 2. **E)** Effect of acquisition rate on detection of exocytic events. Timelapse images of VAMP2-pHluorin taken at 10 Hz had frames removed to artificially create the same time lapse at 5, 2, and 1 Hz. **F)** Examples of simulated images with noise of increasing Gaussian intensity. **G)** Six simulations of exocytic events with increasing Gaussian or Poisson intensity were used to test the robustness of the algorithm to image background and system noise. Typical estimated range of signal-to-background of the data is highlighted in cyan. **H)** Trace of change in fluorescence intensity relative to initial fluorescence (ΔF/F) for an individual exocytic event. Fusion pore opening occurs at t = 0 sec. Several parameters including the normalized peak change in fluorescence intensity (peak ΔF/F), event half-life (t_1/2_, sec) and event shape are measured for each exocytic event. Initial fluorescence is estimated from average of 10 frames (1 sec) prior to highest peak of fluorescence detected (initiation of exocytosis). Half-life is estimated based on fitting a negative exponential decay function to the fluorescence intensity of the tracked event over time. **I)** User-based and automated detection of the frequency of VAMP2-pHluorin-mediated exocytosis in mouse cortical neurons *in vitro* were not different (paired t-test, p = 0.56). Histograms of measured peak ΔF/F **(J)**, event t_1/2_ **(K)**, and major/minor axis of the first frame of each detected exocytic event **(L)**. n = 462 exocytic events for each.

### Confirmation of *bona fide* exocytic detection

To test the algorithm accuracy at identifying transient events, neurons were imaged before and after treatment with NH_4_Cl (Figure 1D, Movie 2), which alkalinizes the pH of intracellular vesicular compartments, reversing the quenching of pHluorin fluorescence (Miesenböck et al., 1998). NH_4_Cl treatment increased vesicular fluorescence in the neurons (Figure 1D) and events detected were near or at zero, indicating the algorithm only identified transient fluorescent exocytic events. To confirm that identified events were dependent upon VAMP2-mediated SNARE complex formation, neurons were treated with Tetanus NeuroToxin (TeNT), which cleaves the cytoplasmic face of VAMP2, preventing SNARE complex formation and fusion (Link et al., 1992). VAMP2-pHluorin-mediated fusion events were rarely detected, if at all, indicating that the algorithm identified TeNT-sensitive events (Figure 1D, Movie 2). To ensure that images were acquired at a sufficient framerate to capture all exocytic events, frames were removed to artificially decrease the framerate. At an acquisition rate of 1 Hz, the frequency of detected events was underestimated as compared to 5 Hz; no further improvement occurred going from 5 Hz to 10 Hz (Figure 1E). Although 10 Hz provided oversampling for estimation of exocytic frequency, we chose this acquisition rate to ensure ample temporal resolution for the analyses of individual exocytic events.

To determine the robustness of the algorithm to local background variation due to plasma membrane localized VAMP2-pHluorin and system noise, and to calculate true error rates, simulations of exocytic events based on empirical measurements were created (Figure 1F). Cell-surface VAMP2-pHluorin signal was simulated as random intensity across cell masks. Gaussian fluorescent peaks with varying means and standard deviations, estimated from hand-counted pHluorin signal, were added to simulate exocytic events. A range of intensities based on typical images that followed a Poisson or Gaussian distribution were added to simulated images to represent random noise (Pawley, 1990; Waters, 2009). The detection software was robust to the typical signal-to-noise ratio observed experimentally, with an error rate between 2-7% (Figure 1G), comparable or better than other algorithms (Sebastian et al., 2006; Yuan et al., 2015).

After identification of events, a series of measurements were made to describe the population of fusing vesicles (Figure 1H-L), including the frequency of exocytosis per neuron (Figure 1I), the peak change in fluorescence for each event (ΔF/F, Figure 1H,J), which estimates the amount of VAMP2-pHluorin per vesicle, the event half-life (t_1/2_, Figure 1H,K), which represents how quickly the VAMP2-pHluorin fluorescence decayed, the shape of the initial exocytic boundary (major/minor axis, Figure 1L), and the time (t) and location (x,y) within the cell mask of fusion. The shape of the half-life distribution exhibited no artificial truncation, indicating that the algorithm captured events with both short and long half-lives. The frequency of VAMP2-pHluorin exocytic events detected by the software was not different from those counted by several independent, trained users (Figure 1I, p=0.56).

### Confirmation of detection of VAMP7-mediated exocytosis

We next tested whether the algorithm, built on VAMP2-mediated events, also accurately detected VAMP7-mediated events. Surprisingly, the frequency of VAMP7-mediated exocytosis detected by the algorithm was considerably lower than we previously reported using non-automated analysis (Figure S1B, Winkle et al., 2014). Alkalization with NH_4_Cl revealed numerous VAMP7-containing vesicles and confirmed VAMP7-pHluorin fluorescence was quenched by low pH, as expected (Fig S3A). To ensure the algorithm was capable of detecting VAMP7-mediated fusion, we co-expressed VAMP7-pHluorin along with an activating VAMP7 mutant lacking the autoinhibitory longin domain (VAMP7ΔLD, Martinez-Arca et al., 2000). VAMP7ΔLD expression increased the frequency of VAMP7-mediated exocytosis (FigS3B), as previously shown (Burgo et al., 2013). This demonstrated the algorithm was capable of detecting VAMP7-mediated events.

Several potential factors could account for the disparity between the algorithm and hand-counted frequencies of VAMP7-mediated exocytosis, including different image acquisition rates, unique attributes of VAMP7-containing vesicles, and the defining features of a fusion event employed by the algorithm compared to more loosely defined user-identified events. Although artificially decreasing the temporal resolution of images to the 2 Hz used in the previous study was associated with a higher exocytic frequency (Figure S3C), this did not fully account for the differences between automated and user-defined values. Since these newly identified events only met the algorithm requirements at reduced temporal resolution, we conclude they were unlikely true exocytic events. We next compared features of VAMP2 and VAMP7-containing objects. The peak ΔF/F and half-life were not significantly different between VAMP2-pHluorin and VAMP7-pHluorin (Figure S3D), indicating these were unlikely the sources of incongruence. Next, we relaxed the algorithm requirement that events be non-motile. Along with artificially lowering the frame-rate to 2 Hz, this change permitted the software to capture an exocytic frequency indistinguishable from previous reports (Figure S3C). Indeed, the mean squared displacement of exocytic events detected under this relaxed requirement was higher for VAMP7 than VAMP2 (Figure S3E), and thus likely explains why VAMP7-mediated exocytic events were previously overestimated, and VAMP2-mediated events were not.

To examine whether motile fluorescent puncta represented actual exocytic events, we co-expressed the autoinhibitory longin domain (VAMP7-LD, Martinez-Arca et al., 2000) with VAMP7-pHluorin. As reported, VAMP7-LD expression decreased detected VAMP7-pHluorin mediated exocytic events, when the non-motile requirement was included in the detection algorithm (FigS3B). Upon relaxation of this parameter, more VAMP7-mediated events were detected (FigS3B), which also showed a high motility (Fig S3E). Together these results suggested that motile fluorescent events were not true exocytic events, and that the algorithm accurately detected exocytosis.

### Spatiotemporal changes in exocytosis occur during neuronal development

With the algorithm validated, we set out to investigate developmental exocytosis. Dissociated neurons initially exhibit a “pancake-like” shape with peripheral protrusions. By 24 hours *in vitro*, protrusions coalesce into immature neurites. By 48 hours, neurons typically have several defined neurites, one of which is significantly longer and considered the axon. By 72 hours, neurons are characterized by longer neurites and a longer, potentially branched axon. We hypothesized that the temporal and spatial distribution of exocytosis in neurons would change over developmental time, and potentially involve fusion of both VAMP2 and VAMP7-containing vesicles. We measured VAMP2-pHluorin and VAMP7-pHluorin exocytic parameters in mouse embryonic (E) day 15.5 cortical neurons after 24 hours, 48 hours, and 72 hours *in vitro* (Figure 2A-I, Movie 3-4). The rate of VAMP7-mediated exocytosis was negligible under these conditions, so we focused on the organization of VAMP2-mediated exocytosis. We mapped the spatial occurrence and density of VAMP2-pHluorin mediated exocytic events onto cellular masks (Figure 2A), which suggested that their spatial distribution was not uniform. Neurons were segmented into soma and neurite regions of interest (ROI), and the frequencies of VAMP2-mediated exocytosis were compared, controlling for ventral cell surface area (Figure 2B-D). At 24 hours *in vitro*, exocytic frequency was faster compared to 48 and 72 hours (Figure 2B), and faster in the soma compared to neurites (Figure 2C-D). The relationship between exocytic frequency in the soma to neurites remained the same between 24 and 48 hours (Figure 2C-D). However, at 72 hours a redistribution occurred, with an increased frequency of events in neurites and decreased frequency in the soma (Figure 2C-D).

**Figure 2.**
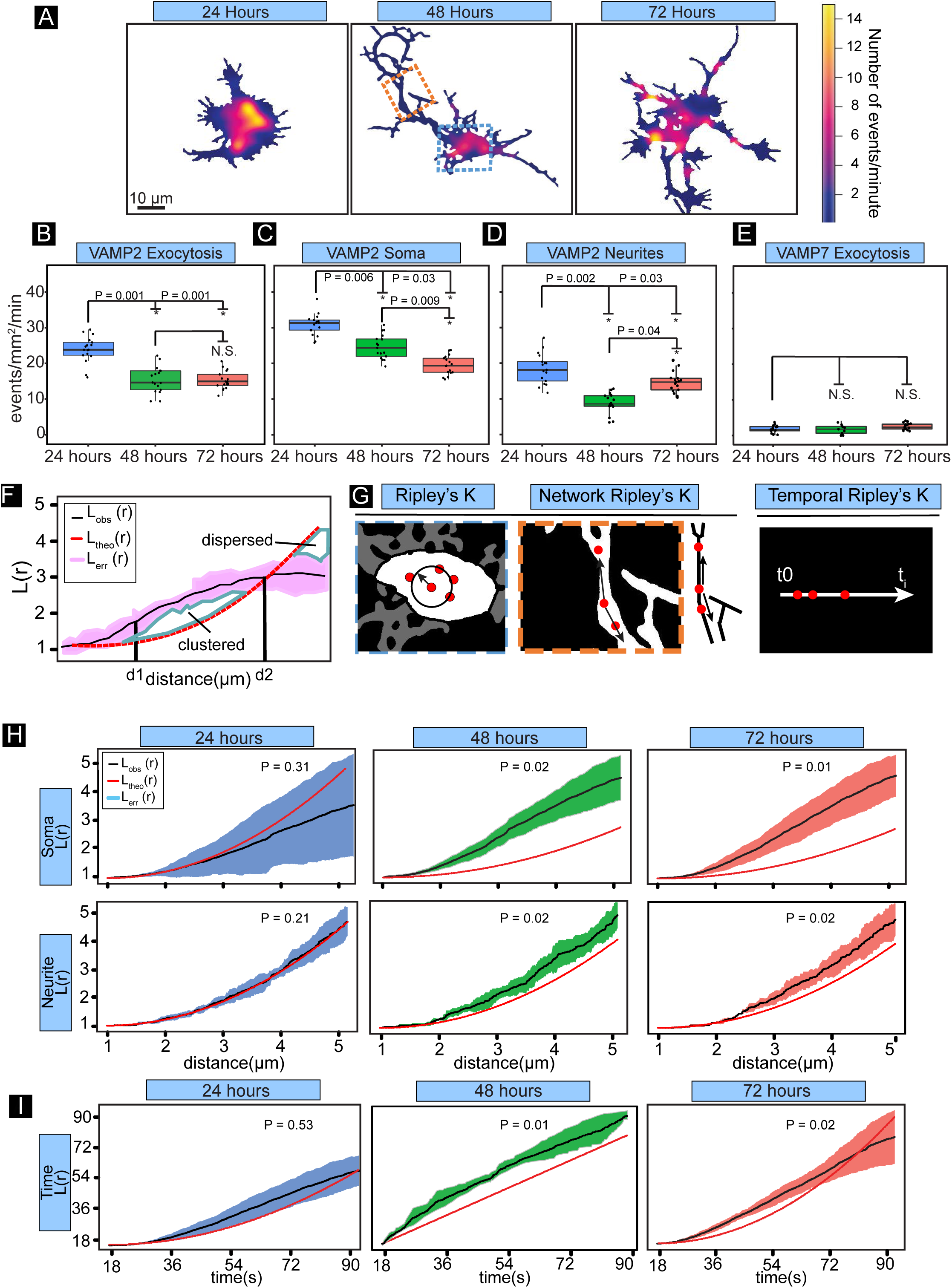
Spatiotemporal changes in exocytosis occur during neuronal development. **A)** Heatmaps of the density of VAMP2-mediated exocytosis in cortical neurons cultured *in vitro* for 24, 48 or 72 hours prior to imaging. **B)** Frequency of VAMP2-pHluorin mediated exocytosis for basal plasma membrane of cortical neurons cultured 24, 48, and 72 hours *in vitro*. (*=P<0.05, n = 17 cells per condition, 464, 455, 466 exocytic events per condition). See Movie 3. **C-D)** Frequency of VAMP2-pHluorin mediated exocytosis in the soma (C) and neurites (D) of cortical neurons cultured 24, 48, and 72 hours *in vitro*. **E)** Frequency of VAMP7-pHluorin mediated exocytosis for basal plasma membrane of cortical neurons cultured 24, 48, 72 hours *in vitro*. The frequency of exocytosis was not significantly different between time points (n = 14 cells per condition). See Movie 4. **F)** Example of Ripley’s L(r) analyses. The mean L(r) value of the aggregated data from multiple replicates (L_obs_(r), black line) and standard error of the data (L_err_(r), pink) is compared to the expected L(r) value of completely random exocytic events (L_theo_(r), red dashed line). **G)** Schematics of Ripley’s L(r) function in both spatial and temporal dimensions. The red dots represent exocytic events. **H)** Spatial Ripley’s L(r) function analysis revealed that events were randomly distributed in the soma at 24 hours, whereas exocytosis occurred in spatial clusters in the soma at 48 hours and 72 hours. Exocytosis followed a similar pattern in the neurites, being randomly distributed in the neurites at 24 hours and clustered at 48 hours and 72 hours. **I)** Temporal Ripley’s L(r) function of exocytic events over time in mouse cortical neurons, yet temporal bursts of exocytosis at 48 and 72 hours *in vitro*. See Movie 3.

### Ripley’s K analysis reveals developmental exocytic clustering

The exocytic heatmaps suggested that events were clustered into spatial hotspots (Figure 2A). To determine if exocytosis was organized non-randomly, we adapted Ripley’s K function, a method for distinguishing between clustering, randomness, and dispersal of points (Ripley et al, 1976), to neuronal exocytosis. Ripley’s K function is defined in two dimensions as

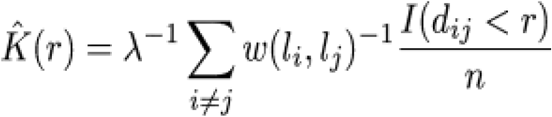

Where λ is the intensity, described as λ = n/A where A is the area encompassing the points, n is the number of points, I is an indicator function (1 if true, 0 if false), d_ij_ is the Euclidian distance between the i^th^ and j^th^ point, and r is the search radius. To correct for edge effects, the function is weighted by w(*l*_i_, *l*_j_), which represents the fraction of search radius that contains A.

The soma of neurons were analyzed using Ripley’s K function. However long, narrow, and branched neurites posed a challenge to the spatial analysis, even with an edge corrector. To accurately measure clustering, we skeletonized neurites, and considered them a 1D network (Figure 2G). Exocytic events in neurites were mapped to the nearest point on the network. We then performed an extension of Ripley’s K function, the network K function, which takes the form

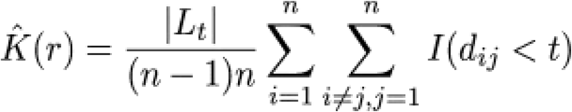
where L_t_ is the union of all links in the network, and d_ij_ is the network distance between points (Okabe and Yamada, 2001).

Finally, to test the temporal clustering of exocytosis, we performed Ripley’s K function in one dimension, which takes on a similar form

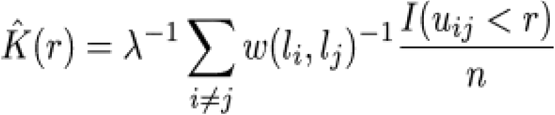

Where u_ij_ is the distance between points on a one-dimensional line (u_ij_ = |u_i_-u_j_|), and w(*l*_i_, *l*_j_) is the fraction of the search length, r, that overlaps the number line (0,T), where T is the length of the one dimensional search space (Figure 2G, total time imaged for temporal analysis).

The K function has been used in a limited setting in exocytosis, performed on individual cells with simple morphology (Díaz et al., 2010, Yuan et al., 2015). To our knowledge, group-comparisons have not been performed on exocytic clustering, which has limited the statistical power and comparisons and conclusions that can be made. With the assumption that all cells measured in a group exhibit the same clustering pattern, the group-specific mean function was defined as

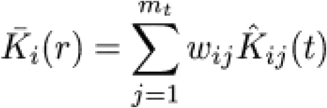

For each i^th^ group, and the overall average K-function mean of the total population is

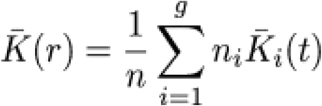

Where w_ij_ = n_ij_/n_i_, n_i_ = the number of points in the i^th^ group, and n = the total number of points of all groups. We constructed a test statistic,

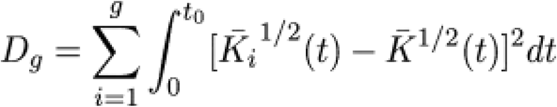

to test differences between groups. This statistic is similar to the residual sum of squares in a conventional one-way ANOVA. We used the average sampling variance of the functions weighted by w_ij_ to construct confidence intervals +/- 2 SE from the mean. To attach P-values to observed test statistics, we used a bootstrapping procedure to determine the distribution of Dg. To do so, we defined

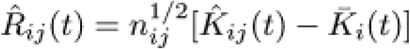

as the residual K functions. We obtained the approximation to the distribution of D_g_ by recomputing its values from the residual K functions by bootstrapping

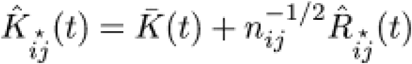
drawing at random with replacement from R_ij_(t), keeping the group sizes fixed. We performed this bootstrapping procedure 999 times, and ranked the observed D_g_ values among the bootstrapped D_g_ values to obtain p-values of how likely our observed values are under the null hypothesis that the two groups being compared came from the same distribution (Diggle, 2013).

For ease of interpretation, we use the variance-stabilized Ripley’s K function, L(r), which gives an L(r) equivalent to the radius (r) under CSR or CTR. To test whether exocytic events were clustered, we compared the L(r) function of aggregated observed data against Monte Carlo simulations of Compete Spatial Randomness (L_CSR_(r)) or Complete Temporal Randomness (L_CTR_(r), Diggle, 2013). Under traditional Ripley’s K analysis, L_CSR_(r) is equivalent to πr^2^; in the case of a network function, L_CSR_(r) is equivalent to the uniform distribution of points on the network. L_CTR_(r) is equivalent to a uniform distribution. At distances where the L_Obs_(r) function rose above spatial or temporal randomness (red line, L_CSR_(r) or L_CTR_(r)), exocytosis was considered clustered, and at distances where the L_Obs_(r) function fell below L_CSR_(r) or L_CTR_(r), exocytic events were considered dispersed (Figure 2F). At 24 hours *in vitro*, exocytosis exhibited spatial randomness in soma and neurites as well as temporal randomness (Figure 2H,I). At 48 and 72 hours *in vitro*, exocytosis was spatially clustered into hotspots in neurites and soma and temporally clustered into bursts (Figure 2H,I). Thus, spatial and temporal organization of exocytosis evolved from random organization to clustered organization during neuronal morphogenesis.

### A mathematical approximation of membrane expansion in developing neurons

To determine the significance of VAMP2-mediated exocytosis, we sought to mathematically calculate neuronal plasma membrane expansion. We estimated the surface area added by VAMP2-mediated exocytosis using the frequency of exocytosis (Figure 2B) and surface area of vesicles. A potential range of vesicular surface areas was approximated from vesicle diameters measured from transmission electron microscopy (TEM) images of cortical neurons at 48 hours *in vitro*, with the assumption that vesicles were spherical (Figure 3A). To correct for capturing random slices through vesicles, we took the conservative estimate that the 25th percent of the interquartile range represented the diameter of the smallest vesicles (80 nm) and the 75th percent of the interquartile-range represented the diameter of largest vesicles (130 nm, Plooster and Menon et al., 2017). We estimated membrane addition between 24 and 48 hours *in vitro*, using the formula

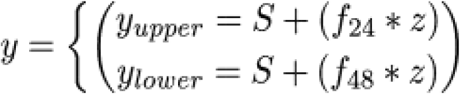

where y represents the predicted neuronal surface area at 48 hours, *S* is the neuronal surface area at 24 hours, *f*_24_ and *f*_48_ are the exocytic frequency at 24 and 48 hours, respectively, and *z* is the surface area of vesicles. Because the frequencies of exocytic events were not measurably different between the basal and lateral membrane (Figure S1), we estimated the total surface membrane of the cells by doubling the apparent area of the basal lateral membrane. The mean frequencies of exocytosis at 24 and 48 hours (Figure 2B) defined upper and lower bounds of VAMP2-mediated membrane addition. The predicted average membrane addition between 24 and 48 hours was ~868-1737 μm^2^ using the smaller vesicle estimation, and ~2200-4587 μm^2^ using the larger vesicle estimation.

**Figure 3:**
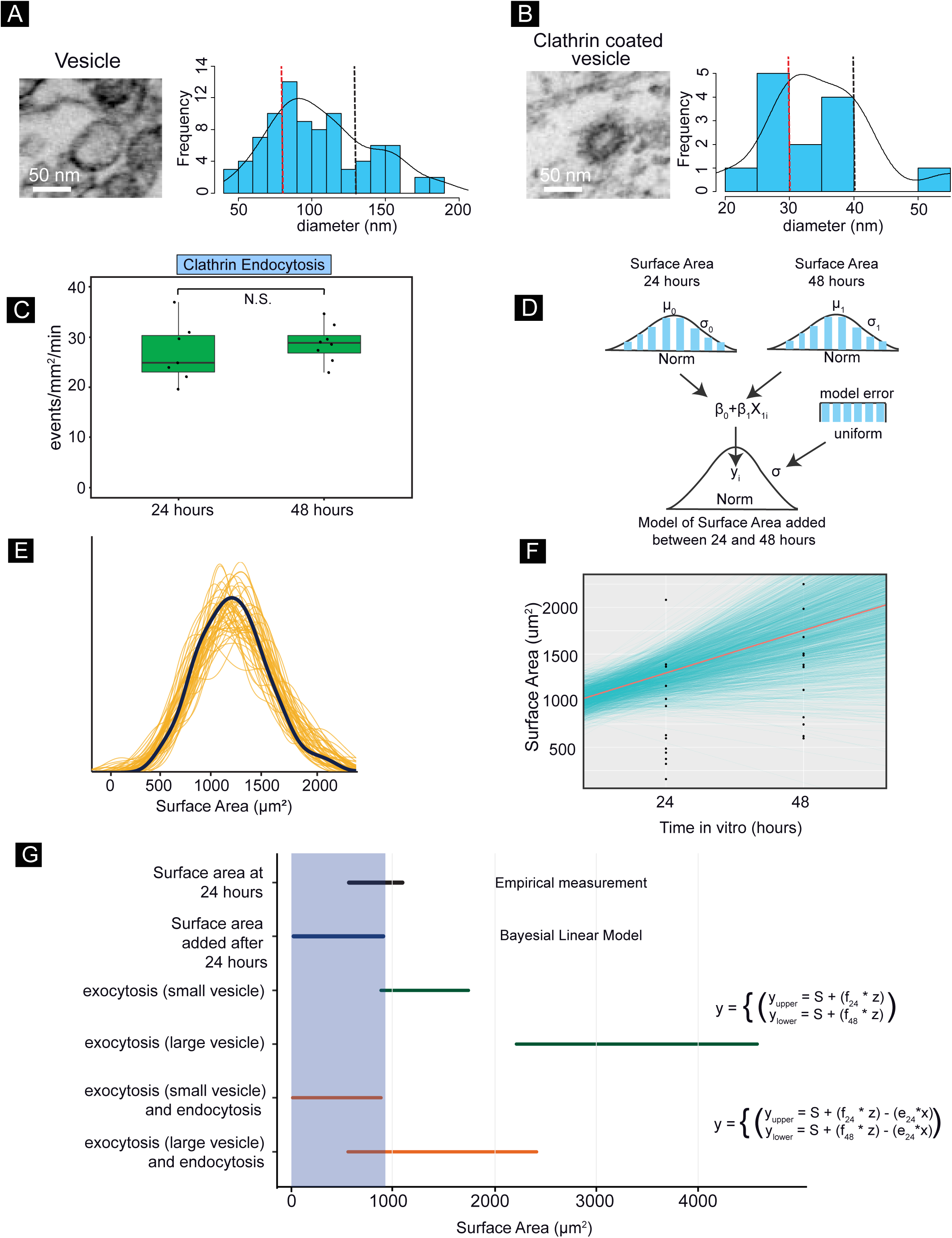
A mathematical estimation of membrane expansion. **A-B)** Example images and histogram of measured diameters of non-clathrin coated vesicle A) and clathrin coated vesicle B) from transmission electron microscopy images of neurons 48 hours *in vitro* with a density line overlaid (black solid line). The 25th percent of the interquartile range is marked with a red dotted line, the 75th percent of the interquartile range is marked with the black dotted vertical line. **C)** Frequency of clathrin-mediated endocytosis at 24 and 48 hours *in vitro* measured from TIRF images of CLC-TagRFP. **D-F)** Hierarchical diagram of the Bayesian linear model and its fit to neuronal surface area increases between 24 and 48 hours. D) Probability distributions (blue) of surface areas at 24 and 48 hours represent priors imposed on the model, with mean μ and standard deviation σ for the normal distributions, and a uniform distribution for the model error (values described in the text). **E)** Posterior predictions of surface area increases by the model (orange lines) were compared to the actual data (black line) to check for model fit. **F)** Credible regression lines (blue) and mean (orange line) regression line of predicted plasma membrane expansion. **G)** Estimates of membrane addition between 24 and 48 hours. The black line is the measured surface area at 24 hours *in* vitro. The blue line and blue shaded area represents the predicted amount of net plasma membrane expansion using a Bayesian linear model (D-F). The green lines represent the predicted amount of membrane added by VAMP2-mediated exocytosis. The orange lines represent the predicted net membrane addition from VAMP2-mediated exocytosis after accounting for membrane removal via clathrin-mediated endocytosis.

We derived the increase in cell surface area between 24 and 48 hours using a Bayesian linear model, based upon empirically measured neuronal surfaces areas, using time *in vitro* as a predictor (Figure 3D). To give more weight to the data, we chose a weakly informative prior distribution for the predictor (normal distribution with mean of 500, sigma of 100), with the assumption that neurons increased surface area over time with no chance of decreasing surface area, and a uniform distribution between 0-10 for the variance. Posterior predictive checking indicated that the model was a good fit (Figure 3E), with predicted credible regression lines showing a positive increase in surface area between 24 and 48 hours *in vitro* (Figure 3F). The 90% confidence interval showed that the surface area increased by only 9-889 μm^2^ with a mean of 447 μm^2^ (Figure 3G, blue line). This amounted to a ~29% increase over a 24-hour time period, in line with previous estimates (Pfenninger, 2009). These results suggested that VAMP2-mediated exocytosis provided excess membrane material for developmental plasma membrane expansion (Figure 3G, green lines).

Excess membrane delivery could be balanced by compensatory endocytosis. Clathrin-mediated endocytosis is conserved from yeast to mammals, and in a number of cell types investigated, 95% of membrane internalization occurs through clathrin-mediated endocytosis, (Bitsikas et al., 2014; Pearse, 1976). Furthermore, ample evidence exists of clathrin-mediated endocytosis at the synapse, yet there is little evidence of clathrin-independent modes of exocytosis in neurons in the literature. We therefore used tagRFP fused to clathrin light chain (Shaner et al., 2008) in conjunction with TIRF microscopy to measure the frequency of clathrin-mediated endocytosis. An h-dome transform was performed to detect fluorescent maxima in each frame. Fluorescent punctae that lasted at least 20 seconds prior to disappearing were considered endocytic events (Aguet et al, 2013). The rate of endocytosis did not change between 24 and 48 hours *in vitro* (Figure 3C). We measured an average diameter of 34.375 nm with an interquartile range of 30-40 nm of clathrin-coated vesicles from TEM images of cortical neurons in culture for 48 hours, which was used to estimate the potential range of surface areas (Figure 3B). With this additional parameter of clathrin-mediated membrane internalization, we updated our formula

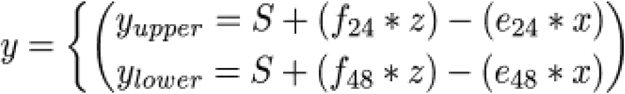

Where e_24_ and e_48_ are the endocytic frequency at 24 and 48 hours and x is the surface area of the clathrin-coated vesicles.

The new predicted average membrane addition between 24 and 48 hours was ~1-868 μm^2^ using the smaller vesicle estimation, and ~547-2416 μm^2^ using the larger vesicle estimation (Figure 3G, orange lines). These results suggest that, by first approximation, plasma membrane expansion in cortical neurons *in vitro* can be explained by VAMP2-mediated exocytosis and clathrin-mediated endocytosis.

### Netrin-1 modulates the spatiotemporal distribution of exocytic events in neurites

To further determine how exocytosis is organized during neuronal morphogenesis, we investigated the consequences of bath application of netrin-1 (Figure 4A-I), a developmental morphogen that accelerates neuronal morphogenesis (Winkle et al., 2016). Further validating the accuracy of the automated software, we confirmed previous findings (Winkle et al., 2014) that netrin-1 stimulation increased the frequency of exocytosis in neurons (Figure 4E). Analysis of individual events identified parameters that were insensitive to netrin-1 stimulation, including the major/minor axis of the initial vesicle fusion event, the peak ΔF/F, and the t_1/2_ of fluorescence decay (Figure 4A-C). This suggests that although netrin-1 altered the probability of vesicle fusion, the same vesicles fuse with the plasma membrane using similar mechanisms, as they contained indistinguishable amounts of VAMP2-pHluorin, and fused and dissipated with identical patterns.

**Figure 4.**
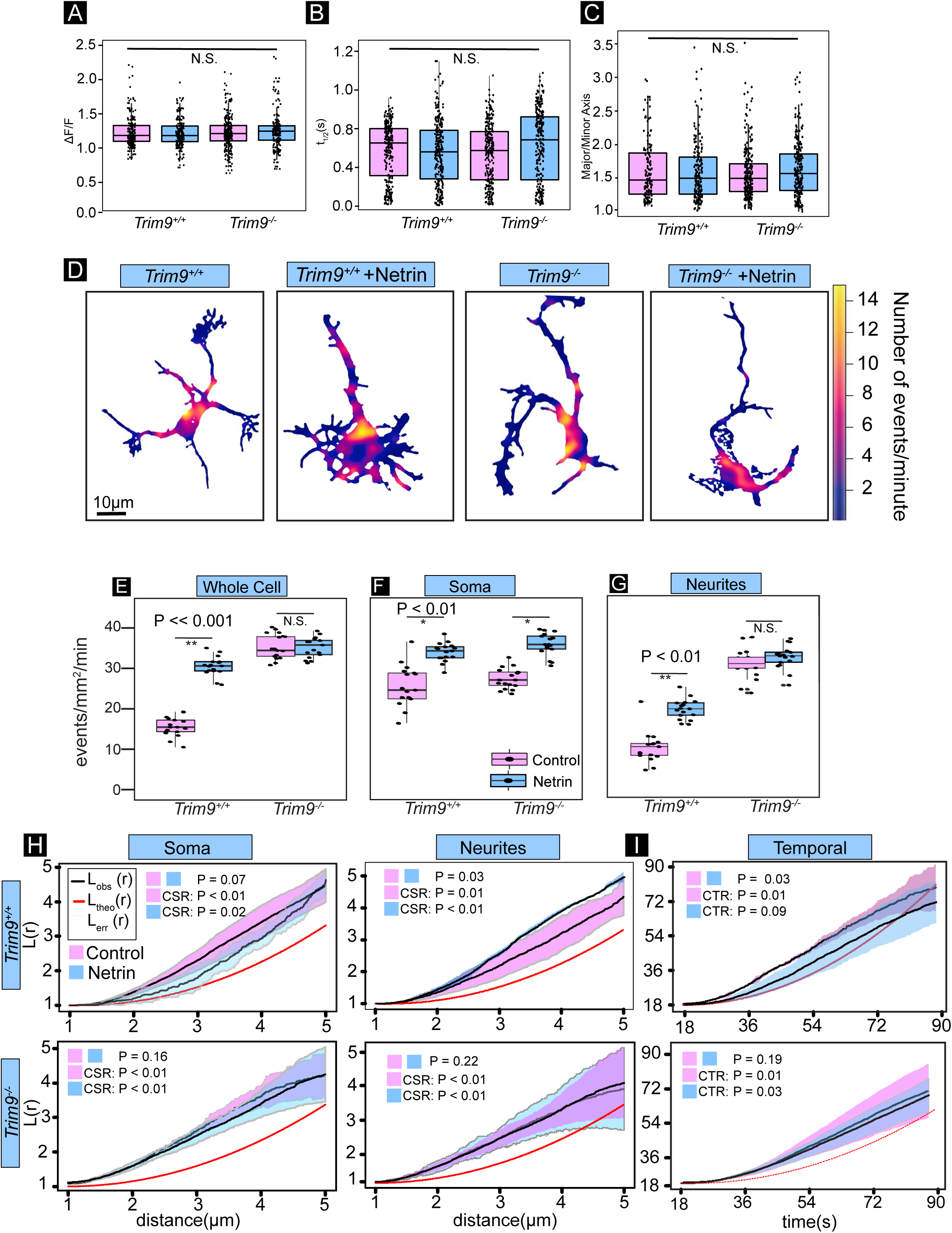
Netrin and TRIM9-dependent changes in the distribution of exocytosis. **A-C)** Individual data points and boxplot of peak ΔF/F (A), t_1/2_ (B), and major/minor axis (C) reveals these parameters of exocytic events were not altered by netrin-1 stimulation or deletion of *Trim9* (n = 562,632,643,673 exocytic events per condition, respectively). **D)** Heatmaps of density of exocytosis in *Trim9*^+*/*+^ and *Trim9*^−/−^ neurons, with and without netrin-1 addition. **E)** Individual data points and boxplot of frequency of VAMP2-pHluorin-mediated exocytosis in *Trim9*^+*/*+^ and *Trim9*^−/−^ cortical neurons, treated or untreated with netrin-1 (boxplot, n = 17 cells per condition). **F-G)** Frequency of VAMP2-pHluorin mediated exocytic events in *Trim9*^+*/*+^ and *Trim9*^−/−^ neurons +/- netrin-1 stimulation in the soma (F) and neurites (G). **H)** Ripley’s L(r) function applied to the soma and neurites of *Trim9*^+/+^ and *Trim9*^−/−^ mouse cortical neurons. Netrin-1 stimulation increased spatial clustering of exocytic events in *Trim9*^+/+^ neurites, but not *Trim9*^-/-^ neurites. Netrin-1 stimulation did not significantly change exocytic clustering in the soma for either the *Trim9*^+/+^ or *Trim9*^−/−^ neurons. **I)** 1D Ripley’s L(r) function of exocytic events over time of *Trim9*^+/+^ and *Trim9*^−/−^ mouse cortical neurons. Exocytosis in *Trim9*^+/+^ neurons was temporally clustered, occurring in temporal bursts over time, which became random upon netrin-1 treatment. In contrast, exocytosis in *Trim9*^−/−^ neurons remained temporally clustered after netrin-1 treatment.

To determine if netrin-1 stimulation changed the spatial or temporal distribution of exocytic events, we mapped the spatial occurrence and density of exocytic events onto cell masks (Figure 4D), which again revealed changes in non-uniform spatial distribution. Interestingly, netrin-1 stimulation increased the frequency of exocytosis in the soma by ~ 31% (Figure 4F), yet increased the frequency of exocytosis in neurites by 92% (Figure 4G), indicating neurites have increased responsiveness to netrin. Ripley’s K function revealed that exocytic events were non-randomly clustered at all distances measured in soma and neurites. In neurites, netrin-treatment enhanced clustering (Figure 4H). In contrast, the spatial clustering in the soma appeared to relax following netrin-1 addition, although these changes were not significant. Ripley’s L(r) function showed temporal exocytic clustering that was lost upon netrin-1 addition (Figure 4I). These results suggest that netrin-1 stimulation modulated the spatial distribution of exocytosis specifically in neurites and dissipated temporal exocytic clustering.

### TRIM9 is required for netrin-dependent exocytic changes in neurites

To determine how netrin differentially affected exocytosis in the soma and neurites, we investigated the brain-enriched E3 ubiquitin ligase TRIM9, previously implicated in constraint of exocytosis and netrin responses (Winkle et al., 2014). Genetic loss of *Trim9* did not change the major/minor axis of the initial vesicle fusion event, the peak ΔF/F, or the t_1/2_ of fluorescence decay (Figure 4A-C). Although *Trim9* deletion increased exocytic frequency and blocked netrin-dependent increases in exocytic frequency (Figure 4E), responses in the soma were similar to *Trim9*^+*/*+^ neurons (Figure 4F), indicating somatic responses were TRIM9-independent. Neurites of *Trim9*^−/−^ neurons however exhibited a ~three-fold increased frequency of basal exocytosis that was not further increased by netrin-1 addition (Figure 4G). This suggests that TRIM9 regulates VAMP2-mediated exocytosis specifically in neurites.

Ripley’s K function indicated that exocytic events were also non-randomly clustered at all distances measured in the soma and neurites of *Trim9*^−/−^ neurons (Figure 4H). Similar to *Trim9*^+*/*+^ neurons, spatial clustering in the soma of *Trim9*^−/−^ neurons appeared to relax following netrin-1 addition (Figure 4H). In contrast, spatial clustering in *Trim9*^−/−^ neurites failed to increase in response to netrin (Figure 4H). Further, Ripley’s L(r) function showed temporal clustering of exocytic events in *Trim9*^-/-^ neurons that persisted following netrin-1 addition (Figure 4I). These results suggested that netrin-1 dependent modulation of the spatial distribution of exocytosis in neurites and the temporal clustering of exocytosis were TRIM9-dependent.

### Different domains of TRIM9 modulate specific parameters of exocytosis in neurites

TRIM9 is a TRIpartite Motif (TRIM) E3 ubiquitin ligase, characterized by an NH_2_-terminal ubiquitin ligase RING domain, two BBox domains that modulate ligase activity, and a coiled-coil (CC) motif that mediates multimerization (Figure 5A). The TRIM9 CC motif also binds the exocytic t-SNARE SNAP25 (Li et al., 2001) and blocks SNARE complex formation and vesicle fusion (Winkle et al., 2014). The TRIM9 COOH-terminal SPRY domain binds the netrin-1 receptor DCC (Winkle et al., 2014). To elucidate how these domains and functions modified the spatial and temporal parameters of exocytosis and netrin-1 responses, we performed structure-function analysis. Expression of full length and domain mutants of TRIM9 (Figure 5A) lacking the RING domain (TRIM9ΔRING), SPRY domain (TRIM9ΔSPRY), or CC motif (TRIM9ΔCC) in *Trim9*^−/−^ neurons further indicated that TRIM9-dependent regulation of exocytic frequency occurred only in neurites (Figure 5B-D). Reintroduction of full length TRIM9 reduced the elevated basal frequency of exocytosis and returned netrin-1 sensitivity to the exocytic frequency in neurites (Figure 5C), the exocytic clustering in neurites (Figure S4A), and relaxation of temporal clustering (Figure S4B).

**Figure 5.**
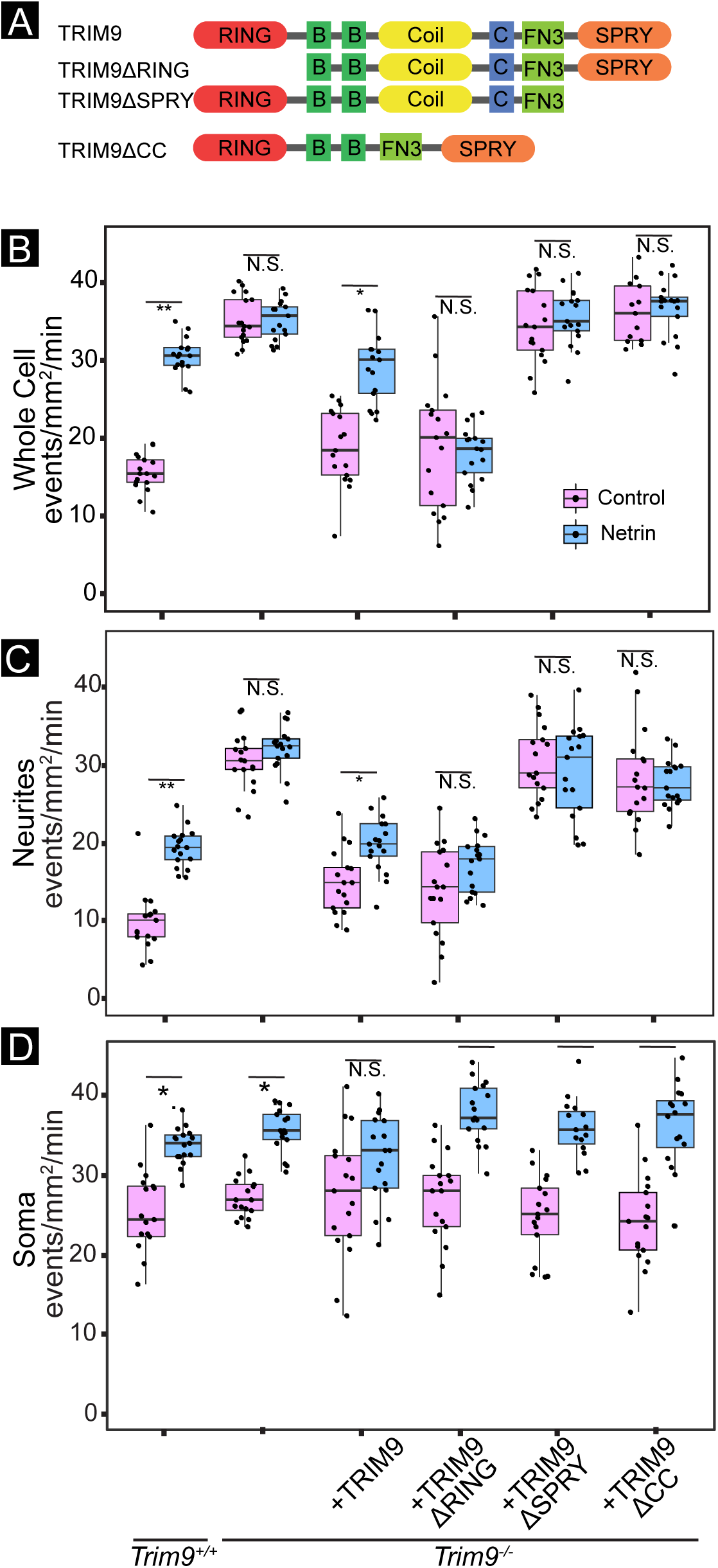
Different domains of TRIM9 modulate specific parameters of exocytosis in neurites. **A)** Domain organization of murine TRIM9 (top) and domain mutants that lack the ubiquitin ligase containing RING domain (TRIM9ΔRING), the DCC binding SPRY domain (TRIM9ΔSPRY), or the coiled-coil motif (TRIM9ΔCC), which mediates TRIM9 multimerization and interaction with SNAP25. **B-D)** Frequency of VAMP2-pHluorin mediated exocytosis across basal plasma membrane of neurons (B), neurites (C), or soma (D) in *Trim9*^+*/*+^ and *Trim9*^−/−^ neurons expressing VAMP2-pHluorin and the indicated TRIM9 domain mutants (boxplots, n = 17 for each condition, * = P < 0.05, ** = P < 0.005). See Figure S4.

TRIM9ΔRING expression reduced the elevated basal frequency of exocytosis in neurites, but failed to rescue netrin-dependent increases in exocytosis (Figure 5C), netrin-dependent spatial clustering of exocytosis (Figure S4A), or netrin-dependent loss of temporal exocytic clustering (Figure S4B). This indicated the ligase domain was unnecessary to constrain exocytosis, but required for response to netrin. Although previous studies identified TRIM9-dependent ubiquitination of substrates as a mechanism to alter substrate function (Menon et al., 2015; Plooster et al., 2017), we did not observe netrin-1 or TRIM9-dependent changes in the ubiquitination of SNAP25 (Figure S4C), suggesting that ubiquitin-dependent modulation of SNAP25 did not explain TRIM9-dependent changes in exocytosis. Expression of TRIM9ΔSPRY or TRIM9ΔCC failed to reduce the elevated frequency of exocytosis in the neurites (P = 0.23 and P = 0.33, respectively, Figure 5C) or the netrin-dependent reduction in temporal clustering (Figure S4B). Expression of TRIM9ΔSPRY, but not TRIM9ΔCC, rescued the spatial exocytic clustering in response to netrin-1 (Figure S4A), suggesting that TRIM9ΔSPRY retains partial netrin sensitivity. Together these results suggested that distinct domains and functions of TRIM9 were specifically required for constraining exocytosis and regulating spatial and temporal clustering in response to netrin-1.

### Exocytosis in interphase melanoma cells is distinctly organized

In contrast to developing neurons, non-neuronal interphase cells are typically at a steady-state size. To determine if spatial and temporal control of exocytosis varied in such a case, we investigated human VMM39 melanoma cells. We monitored exocytosis mediated by two v-SNAREs expressed in melanocytes/melanoma cells, by imaging VAMP3-pHluorin or VAMP7-pHluorin (Figure 6, Movie 5). VAMP3-pHluorin and VAMP7-pHluorin exhibited no difference in the frequency of exocytosis in VMM39 cells (Figure 6A), however this frequency was slower than mediated by VAMP2 and faster than mediated by VAMP7-mediated in neurons (P = 0.002 and 0.017, respectively). Mapping the location and density of VAMP3-and VAMP7-mediated events revealed that spatial distributions of vesicle fusion were distinct (Figure 6B). VAMP7-mediated events appeared polarized, whereas VAMP3-mediated events were distributed throughout the cell. To quantify this distribution, we segmented the cell with lines 10 μm from the edges of the cell, perpendicular to the longest axis (Figure 6B). VAMP7-mediated exocytosis occurred more in the larger end of the cell than in the middle and smaller end, whereas the frequency of VAMP3-mediated exocytosis was similar in each region (Figure 6C).

**Figure 6.**
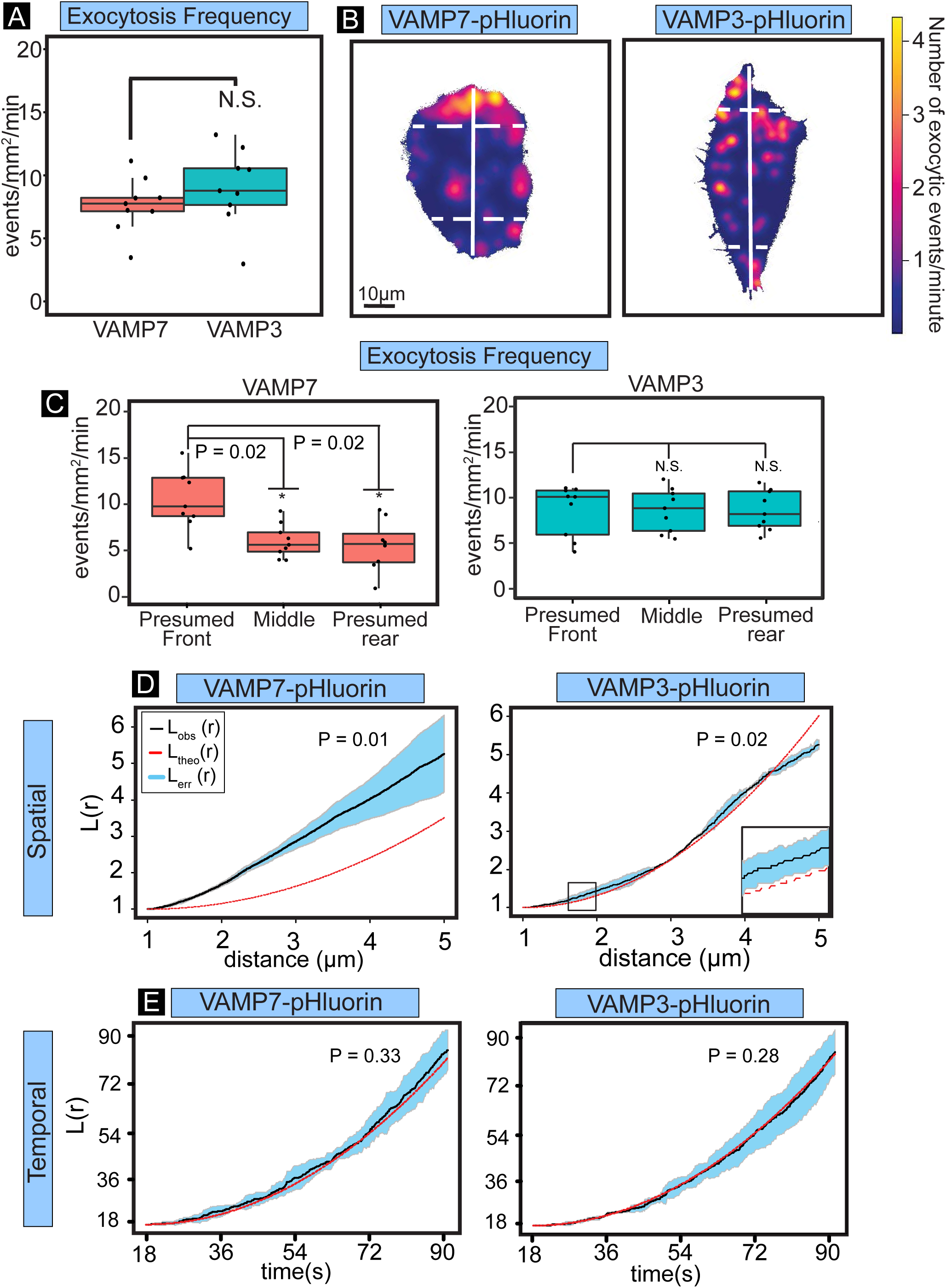
Distinct spatiotemporal organization of fusion of different vesicles pools in interphase melanoma cells. **A)** Frequency of VAMP7-pHluorin and VAMP3-pHluorin mediated exocytosis in VMM39 melanoma cells (boxplots, n = 9 cells/condition). **B)** Heatmaps of density of VAMP7-and VAMP3-mediated exocytosis in representative VMM39 melanoma cells. Solid white line indicates the longest axis of the cell. Dashed perpendicular lines indicate minor axes 10 μm from the tips of the longest axis. The area encompassed from the edge of the cell to the dotted lines were measured to define a larger and smaller end of the cell. **C)** Number of exocytic events normalized to the area in the larger end, middle and smaller end of the cell. VAMP7-mediated exocytosis was polarized to the larger end of the cell. **D)** 2D Ripley’s L(r) function applied to the spatial occurrence of VAMP7-and VAMP3-mediated exocytic events. VAMP7-mediated events exhibited non-random clustering at all distances measured, whereas VAMP3-mediated events were dispersed at larger distances in regularly-spaced clusters 1.5-2.5 μm in size. **E)** 1D Ripley’s L(r) function applied to the temporal occurrence of VAMP7-and VAMP3-mediated exocytic events. Both VAMP7 and VAMP3-mediated events exhibited random temporal occurrence. See Movie 5.

2D Ripley’s L(r) analysis demonstrated that VAMP7-mediated events clustered at all distances measured (Figure 6D). VAMP3-mediated exocytosis was organized distinctly; exocytic clustering was observed at 1.5-2.5 μm, (Figure 6D, inset) however at larger distances (4.5 μm), the Ripley’s L(r) value fell below spatial randomness, indicating dispersal of exocytic hotspots. 1D Ripley’s L(r) analysis of the timing of VAMP3-and VAMP7-mediated exocytic events revealed that both were temporally random (Figure 6E). Together these investigations of the spatial and temporal organization of exocytosis in interphase cells revealed an organization and likely regulation of exocytosis distinct from developing neurons.

## Discussion

To our knowledge this represents the first systematic, automated investigation of exocytosis in developing neurons. Further, although statistics for spatial clustering exist, they have not been applied in cells with extreme polarized morphology nor have they been aggregated to perform comparisons between conditions. Here we built a computer-vision image analysis tool and applied extended clustering statistics to demonstrate that the spatiotemporal distribution of constitutive VAMP2-mediated exocytosis is dynamic in developing neurons, modified by both developmental time and the guidance cue netrin-1, regulated differentially in the soma and neurites, and distinct from exocytosis in non-neuronal cells. Our data and mathematical calculations suggest that in developing neurons, addition of membrane by VAMP2-mediated exocytosis and endocytic retrieval of membrane by clathrin is sufficient to explain, by first approximation, developmental plasma membrane expansion.

### Dynamic spatial regulation of exocytosis in developing neurons

Classic studies hypothesized that vesicles fuse at sites of cell growth (Bergmann et al., 1983; Bretscher, 1996; Craig et al., 1995; Feldman et al., 1981; Pfenninger and Friedman, 1993). Others suggested that membrane translocates anterogradely to the extending neurite tips (Popov et al., 1993), yet a systematic investigation of where vesicle fusion occurred had not been done. Based on our findings, we suggest that at earlier developmental stages, rapid membrane expansion is needed but the localization of insertion is inconsequential. Perhaps with a less complex neuronal shape, cytoskeletal forces are sufficient to shape the neuronal plasma membrane. The developmental reduction in exocytic frequency but emergence of exocytic hotspots and shift of exocytosis toward neurites as neurons acquire a more complex morphology suggests that growth of individual neurites or individual branches may require more local membrane insertion. Differences in the spatial distribution of exocytosis between developmental time points and cellular shapes suggest that the location and clustering of membrane insertion via exocytosis is regulated and developmentally critical to reach appropriate neuronal morphology. Our findings indicate that VAMP2-mediated vesicle fusion occurred proximal to sites of growth, such as at the base of neurites and branches, and that VAMP7-mediated vesicle fusion events were rare at the conditions and developmental time points investigated, potentiall supporting the anterograde membrane flow hypothesis at early developmental time points.

### Calculating membrane growth in developing neurons

Several sources for membrane expansion have been identified, including exocytosis of secretory vesicles and/or lysosomes and membrane transfer at ER-plasma membrane contact sites (Arantes et al., 2006; Pfenninger, 2009; Stefan et al., 2013). Additionally, astrocytes can supply membrane material to neurons through lipoprotein particles (de Chaves et al., 1997; Mauch et al., 2001). The relative contributions of these sources have not been explored. Our results and estimations showed that the predicted neuronal surface area increase supplied by VAMP2-mediated exocytosis was roughly 2-10 times greater than the measured surface area of neurons and that clathrin-mediated endocytosis compensates by removing excess material. VAMP2-mediated exocytosis and clathrin-mediated endocytosis together predict membrane addition that agrees with the membrane growth observed in early developing neurons, suggesting that they may be the predominate mechanisms of membrane flux in developing neurons. However other modes of addition and retrieval are likely involved.

### Exocytosis in the soma and neurites are distinctly regulated during neuronal development

Netrin-1 dependent changes in the spatial constraint of exocytosis only in neurites suggest that VAMP2-containing vesicles exist as two separate populations, or alternatively that the regulatory machinery, such as TRIM9, acts differently in neurites. Different proteins and vesicles have been shown to selectively enter different regions of the developing neuron (Dean et al., 2012; Petersen et al., 2014; Winckler et al., 1999). TRIM9 is present in the soma, yet whether specific TRIM9 isoforms localize and function distinctly, or there is differential regulation of TRIM9 function in the soma and neurites remain to be seen. *In vitro*, we found that cortical axons turn up attractive gradients of netrin-1 (Taylor et al., 2015) and that this is absent in *Trim9*^−/−^ neurons (Menon et al., 2015). Whether a gradient of netrin-1 imparts asymmetrical changes in the spatiotemporal pattern of exocytosis in a TRIM9-dependent fashion has yet to be shown.

### Polarized exocytosis in non-neuronal migrating cells

The slower frequency of exocytosis in melanoma cells as compared to developing neurons is consistent with melanoma cells not actively expanding their membrane, and thus not requiring as many vesicle fusion events. Polarized exocytosis has been suggested as a driving force in directional cell migration (Bretscher, 1996). Indeed, the rate of polarized exocytosis at the leading edge of fibroblasts correlates with the speed of migration, whereas disruption of cell polarization and polarized exocytosis is associated with loss of migration (Bergmann et al., 1983; Bretscher, 1996; Schmoranzer et al., 2003). The role of different VAMPs in migration has not been systematically explored, even though evidence exists of their heterogeneous distribution, transport, and cargo (Advani et al., 1998; Alberts et al., 2006; Bentley and Banker, 2015; Burgo et al., 2012; Oishi et al., 2006). The distinct spatial distributions of VAMP3-and VAMP7-mediated exocytosis observed here suggests unique roles for these two vesicle populations in melanoma cells, with polarized VAMP7-mediated exocytosis potentially associating with expansion of membrane at leading edge during motility.

## Materials and Methods

### Animal Work

All mouse lines were on a C57BL/6J background and bred at UNC with approval from the Institutional Animal Care and Use Committee. Mouse colonies were maintained in specific pathogen-free environment with 12-12 hour light and dark cycles. Timed pregnant females were obtained by placing male and female mice together in a cage overnight; the following day was designated as embryonic day 0.5 (E0.5) if the female had a vaginal plug. *Trim9*^−/−^ mice were described in Winkle et al., 2014. Mice were genotyped using standard tail genotyping procedures.

### Media, culture and transfection of primary neurons and immortalized cell lines

Both male and female embryos were used to generate cultures and were randomly allocated to experimental groups. Cortical neuron cultures were prepared from day E15.5 embryos as previously described (Viesselmann et al., 2011). Briefly, cortices were microdissected and neurons were dissociated with trypsin and plated on poly-D-lysine coated glass-bottom culture dishes in neurobasal media supplemented with B27 (Invitrogen). This same media was used for all time-lapse experiments. For transfection, neurons were resuspended after dissociation in solution (Amaxa Nucleofector; Lonza) and electroporated with a nucleofector according to manufacturer’s protocol. Recombinant netrin was concentrated from the supernatant of HEK293 cells (Serafini et al., 1994). Neurons were stimulated with 500 ng/ml of netrin-1 added 1 hour before imaging.

CRISPR generated *Trim9*^−/−^ HEK293 cells were described previously (Menon et al., 2015). HEK293 and VMM39 cells were maintained in DMEM + glutamine supplemented with 10% FBS. For HEK293 and VMM39 cell transfections, HEK293 cells were transfected using Lipofectamine 2000 as per manufacturer protocol. A list of recombinant DNA used in this paper are listed in Supplementary Table 1.

### Live imaging and image analysis

All time-lapse imaging was performed using an inverted microscope (IX81-ZDC2) with MetaMorph acquisition software, an electron-multiplying charge-coupled device (iXon), and a live cell imaging chamber (Tokai Hit Stage Top Incubator INUG2-FSX). The live cell imaging chamber maintained humidity, temperature at 37°C, and CO_2_ at 5%.

For all primary neuron exocytosis assays, neurons expressing VAMP2-pHluorin or VAMP7-pHluorin were imaged at the indicated timepoint *in vitro* with a 100x 1.49 NA TIRF objective and a solid-state 491-nm laser illumination at 35% power at 110-nm penetration depth or in widefield epifluorescence using Xenon lamp. Images were acquired using stream acquisition, imaging once every 100ms for 2 minutes with 100 ms exposure time (10 Hz) or indicated acquisition rate. For melanoma (VMM39) cell line exocytosis assays, images of interphase cells were acquired at 24 hours after transfection with the indicated VAMP-pHluorin construct with a 100x 1.49 NA TIRF objective and a solid-state 491-nm laser illumination at 35% power at 110-nm penetration depth. Images were acquired every 500 ms for 3 minutes. For all primary neuron endocytosis assays, neurons expressing clathrin light chain-tagRFP were imaged at the one frame per second *in vitro* with a 100x 1.49 NA TIRF objective and a solid-state 561-nm laser illumination at 40% power at 110-nm penetration depth. ImageJ software was used for general processing of images and movies.

### NH_4_CI and TeNT treatments

For these assay, neurons were transfected with VAMP2-pHluorin or VAMP7-pHluorin and imaged at 48 hours. Neurons were imaged as above. Either 25 mM of NH_4_Cl or 50 nM of TeNT (Sigma) were added to the neurons followed by either immediate imaging for NH_4_Cl treatment or imaging 30 minutes after the addition of TeNT. Analysis of exocytic events were performed as described below.

### Ubiquitination assays

To measure ubiquitination, HEK293 cells were transfected with HA-DCC, FLAG-ubiquitin, and SNAP25-GFP using Lipofectamine 2000 (Invitrogen) and cultured for 24 hours. 4 hours prior to lysing, the cells were treated with 10 μM MG-132 or 10 μM MG-132 and 600 ng/ml of netrin-1. The treated HEK cells were lysed in 20 mM Tris-Cl, 250 mM NaCl, 3 mM EDTA, 3 mM EGTA, 0.5% NP-40, 1% SDS, 2mM DTT, 5 mM NEM (N-ethylmaleimide), 3mM iodoacetamide, protease and phosphatase inhibitors, pH = 7.4. For 6 million cells, 270 μl of lysis buffer was added and incubated on ice for 10 minutes. Cells were removed from the dish and transferred into tubes. 30 μl of 1x PBS was added and samples were gently vortexed. Samples were boiled immediately for 20 minutes until they turned clear, then were centrifuged at 14,000 rcf for 10 minutes. The boiled samples were diluted using lysis buffer without SDS to reduce the SDS concentration to 0.1%. For immunoprecipitation of GFP-SNAP25, IgG-conjugated A/G beads (sc-2343, SCBT) were utilized to preclear lysates for 1.5 hours at 4°C with agitation. A/G beads coupled to anti-GFP Ab (Neuromab) were agitated within pre-cleared lysates overnight at 4°C to precipitate GFP-SNAP25. Beads were washed three times with lysis buffer and bound proteins were resolved by SDS-PAGE and analyzed by immunoblotting.

### Statistical analysis

The software package R was primarily used for statistical analysis of data. Both R and Adobe Illustrator were used for the generation of figures. At least three independent experiments were performed for each assay. Normality of data was determined by Shapiro-Wilkes test. For two-sample comparisons of normally distributed data, an unpaired t-test was used for two independent samples, or paired t-test for paired samples. For multiple comparisons, ANOVA was used to determine significance, followed by unpaired or paired t-tests corrected using bonferroni-holms method. For analysis of non-normally distributed data, the Mann-Whitney test was used for two samples. For more than two samples, Kruskal-Wallis non-parametric ANOVA was used to determine significance followed by the method described above. For all Ripley’s L(r) functions, comparisons between individual L(r) functions were made using the studentized permutation test (Hahn, 2012). To test for differences between groups of L(r) functions, we first aggregated K functions and performed Diggle’s test, described in the text (Diggle et al., 1991).

### Ripley’s K test of complete spatial randomness or complete temporal randomness

We performed Monte Carlo simulations of complete spatial or temporal randomness to determine whether each aggregate Ripley’s K function comes from a random distribution (Diggle, 2013). In the case of a 2D Ripley’s K function, complete spatial randomness, or K_csr_(r), is equivalent to πr^2^. As all of our time intervals for exocytosis were equal, we normalized all time data to the interval [0,1]. For a uniform distribution on the interval [0,10], LCTR(r) is equivalent to 1-(1-d)^2^. Tests of spatial or temporal randomness were performed as described in Diggle, 2013. In the case of a network function, L_CSR_(r) is equivalent to the uniform distribution of points on the network.

## Acknowledgments

We thank Patrick Brennwald and the Gupton laboratory for thoughtful critique of this work. We also thank Caroline Monkiewicz, Charles Park, Divya Mahesh, and Emily Wolfgram for their contributions and technical support. We thank the Computational Image Analysis course at the Marine Biological Laboratory at Woods Hole for providing the foundation and expertise for building the image analysis software. We thank the Initiative for Maximizing Student Diversity for their continued support. This work was supported by NIH grant GM108970 (S.L. Gupton) and NS103586 (F.L.Urbina).

---

**Figure S1.**
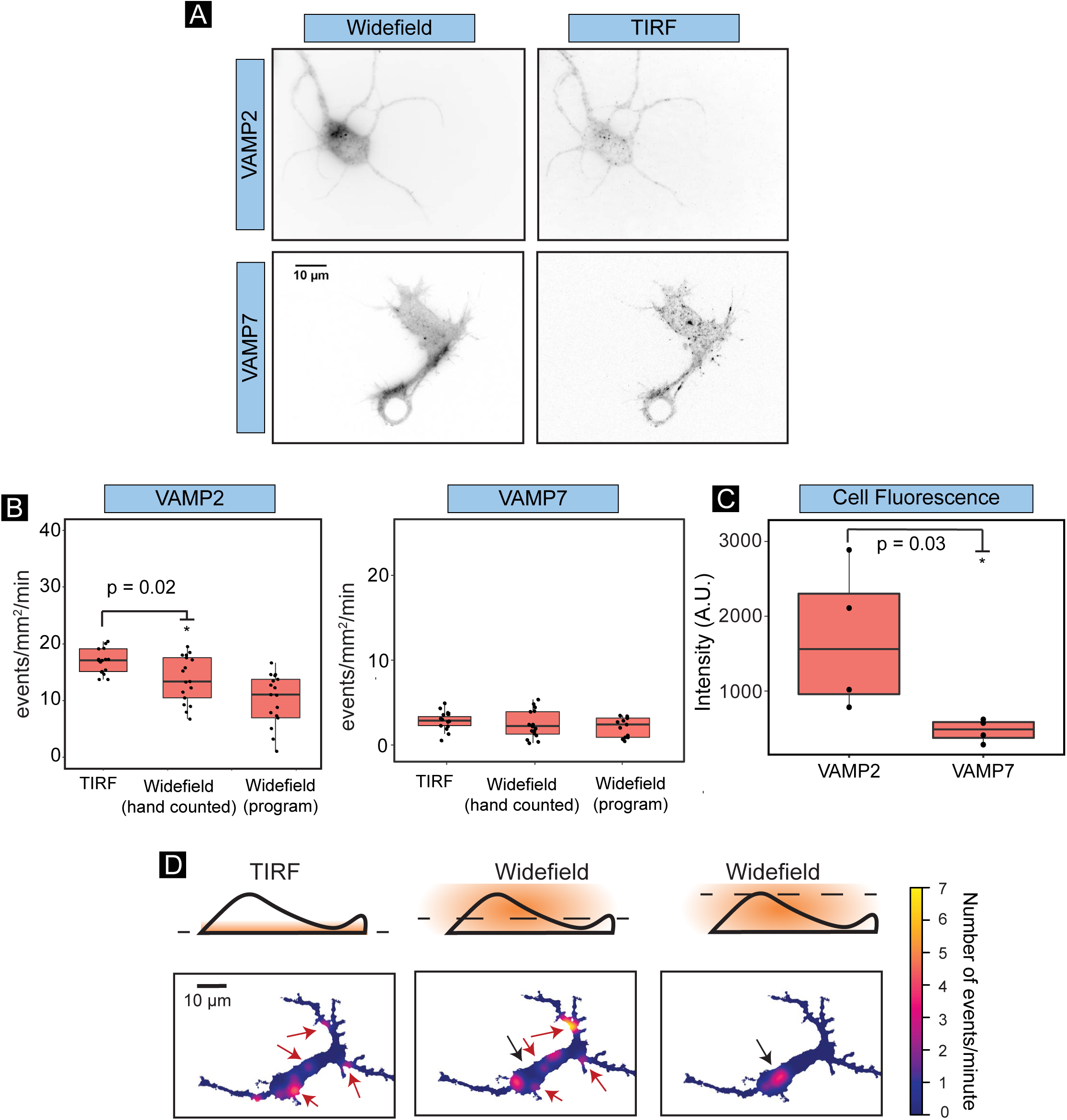
Widefield and TIRF imaging of VAMP2-pHluorin and VAMP7-pHluorin. **A)** Single images from widefield and TIRF timelapses of cortical neurons expressing either VAMP2-pHluorin or VAMP7-pHluorin 48 hours *in vitro*. **B)** Frequency of VAMP2-pHluorin and VAMP7-pHluorin exocytosis measured from TIRF and widefield imaging. **C)** Background plasma membrane fluorescence levels measured from widefield images of VAMP2-pHluorin and VAMP7-pHluorin expressing neurons. **D)** Heatmaps of density of VAMP2-pHluorin exocytosis observed by TIRF (left) and widefield microscopy at bottom (middle) and top (right) focal planes. Black arrows demarcate how apparent exocytic frequency detection changed with the focal plane. Pink arrows denote clusters apparent by TIRF and at the lower focal plane in widefield, that were absent in the higher focal plane in widefield images.

**Figure S2.**
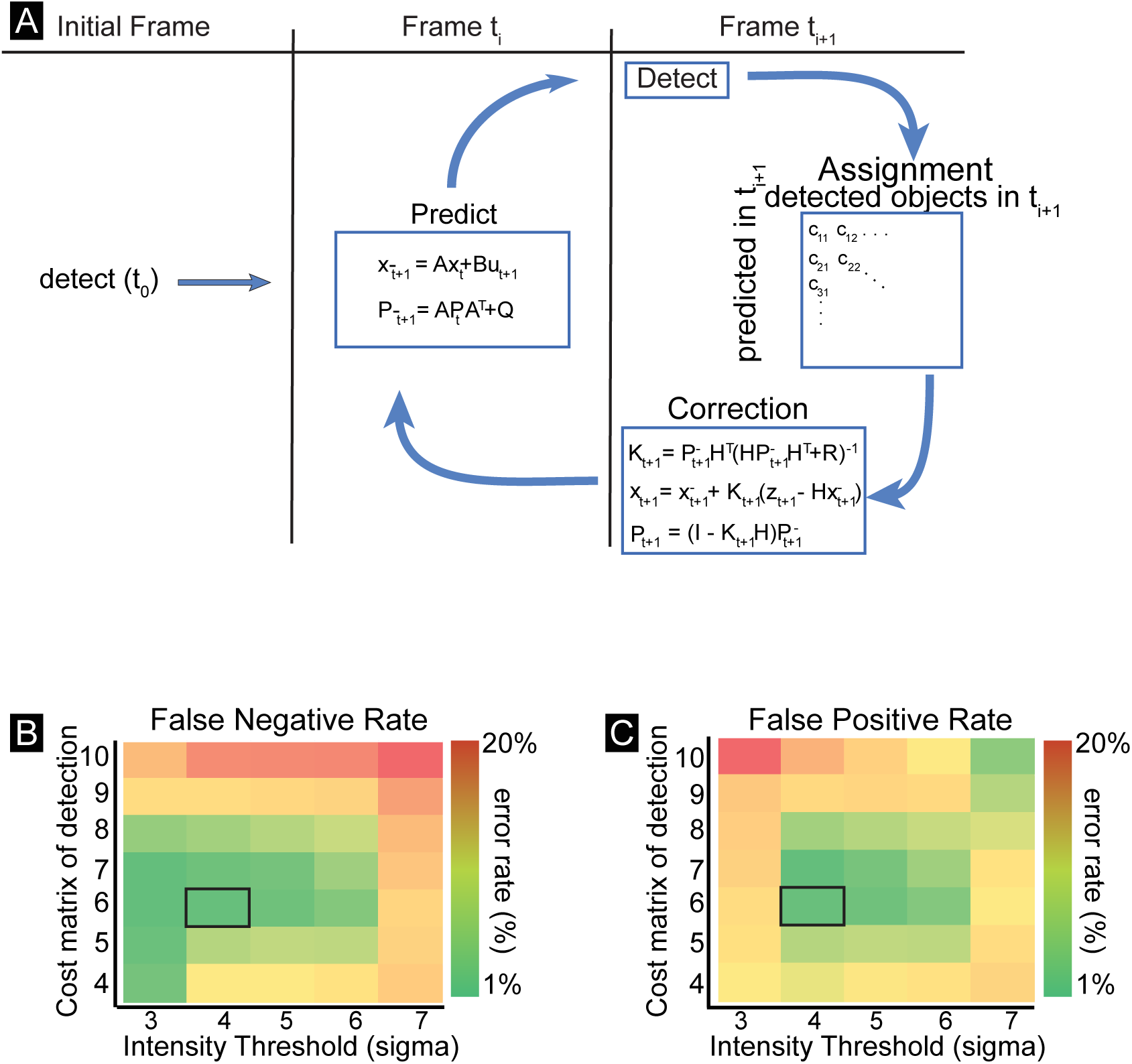
Kalman filter algorithm and parameter selection. **A)** Schematic of the Kalman filter used for linking potential exocytic events between frames. Fluorescent objects were detected in the first frame (t_0_). Using a prediction function, which took into account velocity and current location of the detected fluorescent object, the location of each object was predicted for the next frame (t_+1_). The prediction function was initialized with an experimentally determined set of parameters. After the prediction was made, fluorescent objects were detected in the next frame (t_+1_). These detected objects were then assigned as either a new object, or as a previously detected object based on a cost matrix, which assigned a cost between the distance of predicted objects and detected objects. The penalties assigned to this cost matrix were determined experimentally (B,C). After the newly detected objects were assigned, the prediction function was corrected based upon the error between predicted object locations and detected object locations. The cycle was then repeated, predicting the next location of objects in frame t+2. Cost matrices of the false negative **(B)** and false positive **(C)** rates of event detections with using varying intensity thresholds for detecting events. This indicates how high the signal-to-background ratio must be for pixels to be recognized as a potential exocytic event. False positives rates were computed as 100*(1-M/D) and false negative rates as 100*(1-M/G), where D is the number of computer-detected exocytic events, G is the number of user-defined detected exocytic events, and M are the number of matches between the two, as described in (Matov et al., 2010).

**Figure S3:**
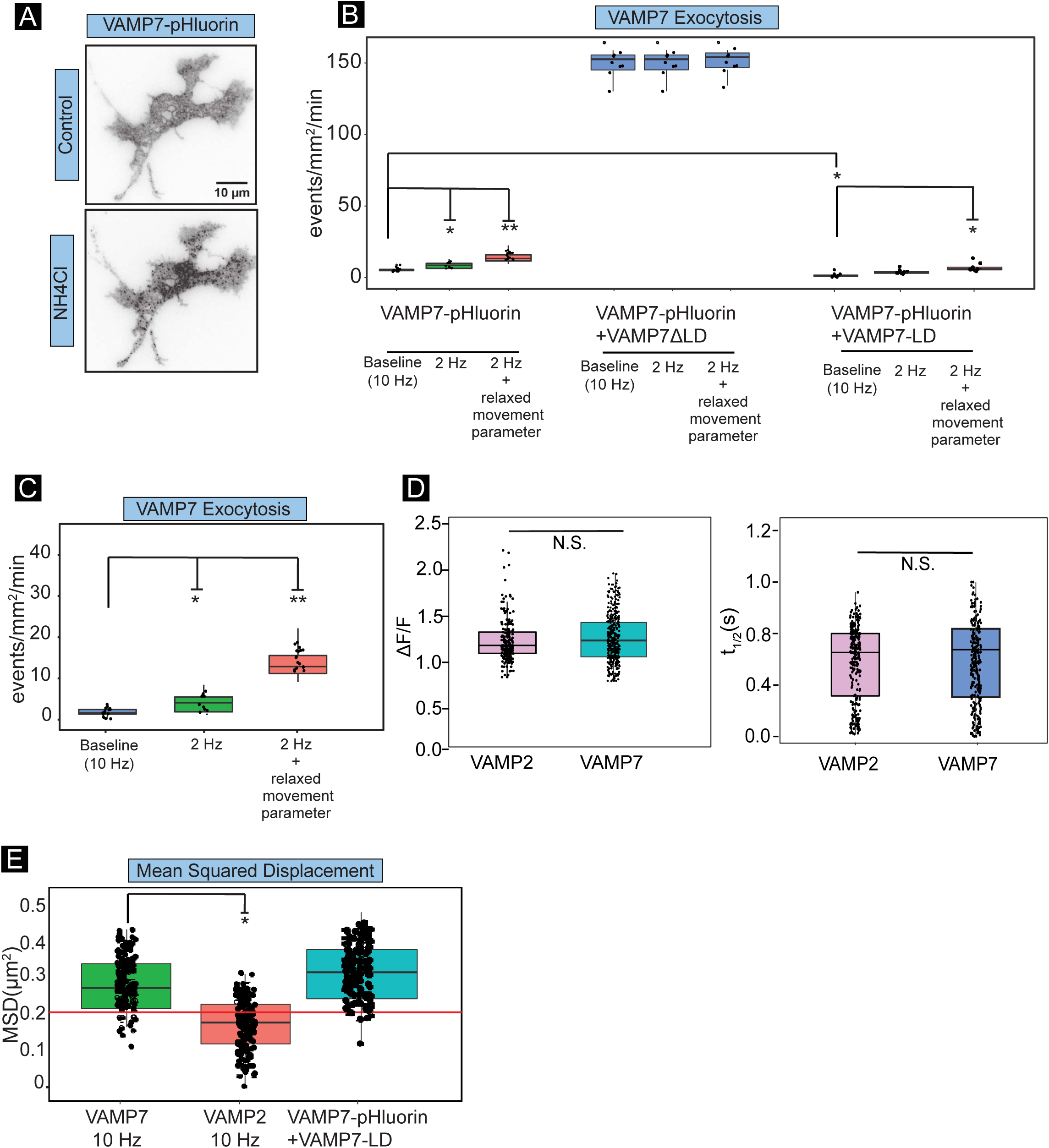
Confirmation of *bona fide* VAMP7-pHluorin exocytosis. **A)** Example minimal projections of TIRF timelapse images of VAMP7-pHluorin before and after treatment with NH_4_CI. **B)** The detected frequency of exocytosis of VAMP7-pHluorin, VAMP7-pHluorin + VAMP7ΔLD, or VAMP7-pHluorin + VAMP7-LD at different acquisition rates and different movement parameter thresholds. **C)** The detected frequency of VAMP7-pHluorin exocytosis at baseline (10 Hz), at 2 Hz, and at 2 Hz with the movement parameter threshold of detection relaxed (p = 0.02 between baseline and 2 Hz, p = 0.001 between baseline and 2 Hz + relaxed movement parameter). **D)** Change in peak fluorescence and half-life of VAMP2-pHluorin and VAMP7-pHluorin mediated events. **E)**. The Mean Squared Displacement (MSD) of potential exocytic events. The red line indicates the experimentally-produced threshold, based on the diffraction-limited resolution of the microscope.

**Figure S4:**
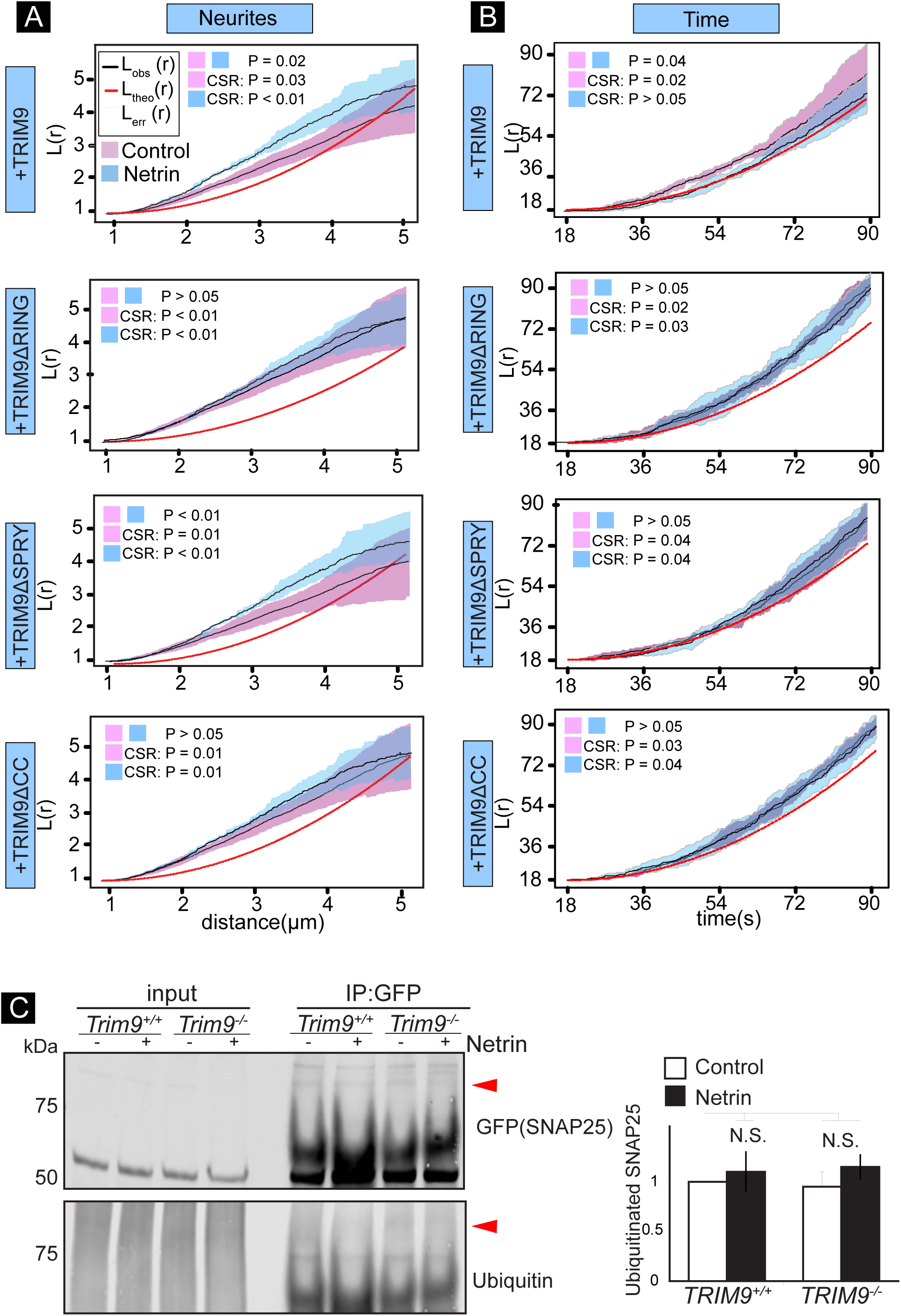
Specific domains of TRIM9 modulate distinct parameters of exocytosis in neurites. **A-B)** 1D Ripley’s L(r) function of spatial distribution in neurites (A) and temporal distribution (B) of exocytosis of *Trim9*^−/−^ mouse cortical neurons expressing the indicated TRIM9 domain mutant. Expression of TRIM9 rescued netrin-1 dependent spatial clustering in neurites and netrin-1 dependent temporal relaxation of exocytic clustering lost in *Trim9*^−/−^ neurons. The RING and CC domain of TRIM9 were required to rescue the netrin-dependent spatial clustering. **C)** SDS-PAGE and western blot of inputs and GFP-SNAP25 immunoprecipitates from *Trim9*^+*/*+^ and *Trim9*^−/−^ HEK293 cells immunoblotted for GFP and ubiquitin. High molecular weight ubiquitin in the IP is considered SNAP25 ubiquitination (arrowheads). Plot shows quantification of SNAP25 ubiquitination. Ubiquitination of SNAP25 did not change significantly in the absence of *TRIM9* or the presence of netrin-1. P-values calculated from Kruskal Wallis ANOVA followed by LSD post-hoc correction compared to *Trim9*^+*/*+^ control.

## Movie Legends

### Movie 1: Identification of exocytic events

(**Left**) Inverted contrast images of E15.5 cortical neuron cultured for 48 hours *in vitro* expressing VAMP2-pHluorin were acquired every 100 ms by TIRF microscopy with 110 nm penetration depth to visualize VAMP2-mediated exocytosis, which is marked by a rapid increase, diffusion and decrease in fluorescence intensity. (**Right**) Same time-lapse is overlaid with detected fluorescent events that meet the criteria of exocytosis events (red): non-motile, transient (<20 frames), diffraction-limited fluorescent events that reach peak fluorescence in the first frame (t_0_) and decay over subsequent frames (t_+i_), and detected events that fail to reach these criteria (blue).

### Movie 2: Confirmation that detected exocytic events are *bona fide*

Inverted contrast images of E15.5 cortical neuron cultured for 48 hours *in vitro* expressing VAMP2-pHluorin were acquired every 100 ms by TIRF microscopy with 110 nm penetration depth to visualize VAMP2-mediated exocytosis, which is marked by a rapid increase, diffusion and decrease in fluorescence intensity. (**Left**) Treatment with NH_4_CI alkalinizes intracellular compartments, reversing the quenching of VAMP2 pHluorin and making VAMP2-pHluorin containing vesicles fluorescent prior to fusion pore opening. (**Right**) Tetanus Neurotoxin (TeNT) treatment, which cleaves the cytoplasmic face of VAMP2, blocks SNARE complex and VAMP2-mediated exocytosis.

### Movie 3: VAMP2-mediated exocytosis changes over developmental time

Inverted contrast images of E15.5 cortical neurons cultured for 24 (left), 48 (middle) and 72 (right) hours *in vitro* expressing VAMP2-pHluorin were acquired every 100 ms by TIRF microscopy with 110 nm penetration depth to visualize VAMP2-mediated exocytosis.

### Movie 4: VAMP7-mediated exocytosis adds negligible amount of membrane

Inverted contrast images of E15.5 cortical neurons cultured for 48 hours *in vitro* expressing VAMP7-pHluorin were acquired every 100 ms by TIRF microscopy with 110 nm penetration depth to visualize VAMP2-mediated exocytosis.

### Movie 5: VAMP3 and VAMP7-mediated exocytosis in VMM39 melanoma cells

Inverted contrast images of interphase VMM39 melanoma cells expressing VAMP3-pHluorin (Left) and VAMP7-pHluorin (right) were acquired every 500 ms by TIRF microscopy with 110 nm penetration depth to visualize VAMP-mediated exocytosis, which is marked by a rapid increase, diffusion and decrease in fluorescence intensity.

## Bibliography

Aalto, M.K., Ronne, H., and Keränen, S. (1993). Yeast syntaxins Sso1p and Sso2p belong to a family of related membrane proteins that function in vesicular transport. EMBO J. 12, 4095–4104.

Advani, R.J., Bae, H.R., Bock, J.B., Chao, D.S., Doung, Y.C., Prekeris, R., Yoo, J.S., and Scheller, R.H. (1998). Seven novel mammalian SNARE proteins localize to distinct membrane compartments. J. Biol. Chem. 273, 10317–10324.

Aguet, F., Antonescu, C.N., Mettlen, M., Schmid, S.L., and Danuser, G. (2013). Advances in analysis of low signal-to-noise images link dynamin and AP2 to the functions of an endocytic checkpoint. Dev. Cell 26, 279–291.

Alberts, P., Rudge, R., Irinopoulou, T., Danglot, L., Gauthier-Rouvière, C., and Galli, T. (2006). Cdc42 and Actin Control Polarized Expression of TI-VAMP Vesicles to Neuronal Growth Cones and Their Fusion with the Plasma Membrane. Mol. Biol. Cell 17, 1194–1203.

Arantes, R.M.E., and Andrews, N.W. (2006). A role for synaptotagmin VII-regulated exocytosis of lysosomes in neurite outgrowth from primary sympathetic neurons. J. Neurosci. 26, 4630–7.

Bentley, M., and Banker, G. (2015). A Novel Assay to Identify the Trafficking Proteins that Bind to Specific Vesicle Populations. Current Protocols in Cell Biology. pp. 13.8.1–13.8.12.

Bergmann, J.E., Kupfer, A., and Singer, S.J., (1983). Membrane insertion at the leading edge of motile fibroblasts. Proc. Natl. Acad. Sci. U. S. A. 80, 1367–1371.

Bishop, C.M. (2006). Mixture Models and EM. In Pattern Recognition and Machine Learning (Springer Science+Business Media, LCC), pp. 423–455.

Bitsikas, V., Corrêa, I.R., and Nichols, B.J. (2014). Clathrin-independent pathways do not contribute significantly to endocytic flux. Elife 3, p. e03970.

Bloom, M., Evans, E.,and Mouritsen, O.G. (1991). Physical properties of the fluid lipid-bilayer component of cell membranes: a perspective. Q. Rev. Biophys. 24, 293–297.

Bowser, D.N., and Khakh, B.S. (2007). Two forms of single-vesicle astrocyte exocytosis imaged with total internal reflection fluorescence microscopy. Proceedings of the National Academy of Sciences 104, 4212–4217.

Brennwald, P., Kearns, B., Champion, K., Keränen, S., Bankaitis, V., Novick, P., Rothman, J.E., Larhammar, D., and Jahn, R. (1994). Sec9 is a SNAP-25-like component of a yeast SNARE complex that may be the effector of Sec4 function in exocytosis. Cell 79, 245–58.

Bretscher, M.S. (1996). Moving membrane up to the front of migrating cells. Cell 85, 465–467.

Burgo, A., Proux-Gillardeaux, V., Sotirakis, E., Bun, P., Casano, A., Verraes, A., Liem, R.K.H., Formstecher, E., Coppey-Moisan, M., and Galli, T. (2012). A molecular network for the transport of the TI-VAMP/VAMP7 vesicles from cell center to periphery. Dev. Cell 23, 166–180.

Burgo, A., Casano, A.M., Kuster, A., Arold, S.T., Wang, G., Nola, S., Verraes, A., Dingli, F., Loew, D., Galli, T. (2013). Increased activity of the vesicular soluble N-ethylmaleimide-sensitive factor attachment protein receptor TI-VAMP/VAMP7 by tyrosine phosphorylation in the Longin domain. J. Biol. Chem. 288, 11960–11972.

Craig, A.M., Wyborski, R.J., and Banker, G. (1995). Preferential addition of newly synthesized membrane protein at axonal growth cones. Nature 375, 592–594.

Dean, C., Liu, H., Staudt, T., Stahlberg, M.A., Vingill, S., Bückers, J., Kamin, D., Engelhardt, J., Jackson, M.B., Hell, S.W., and Chapman, E.R. (2012). Distinct subsets of Syt-IV/BDNF vesicles are sorted to axons versus dendrites and recruited to synapses by activity. J. Neurosci. 32, 5398–5413.

de Chaves, E.I., Rusiñol, A.E., Vance, D.E., Campenot, R.B., and Vance, J.E. (1997). Role of lipoproteins in the delivery of lipids to axons during axonal regeneration. J. Biol. Chem. 272, 30766–30773.

Díaz, E., Ayala, G., Díaz, M.E., Gong, L.-W., and Toomre, D. (2010). Automatic detection of large dense-core vesicles in secretory cells and statistical analysis of their intracellular distribution. IEEE/ACM Trans. Comput. Biol. Bioinform. 7, 2–11.

Diggle, P.J., (2013). Statistical Analysis of Spatial and Spatio-Temporal Point Patterns, Third Edition. CRC Press.

Diggle, P.J., Lange, N., Benes, F.M., (1991). Analysis of Variance for Replicated Spatial Point Patterns in Clinical Neuroanatomy. J. Am. Stat. Assoc. 86, 618.

Engle, and E.C. (2010). Human Genetic Disorders of Axon Guidance. Cold Spring Harb. Perspect. Biol. 2, a001784–a001784.

Feldman, E.L., Axelrod, D., Schwartz, M., Heacock, A.M., and Agranoff, B.W. (1981). Studies on the localization of newly added membrane in growing neurites. J. Neurobiol. 12, 591–598.

Galli, T., Zahraoui, A., Vaidyanathan, V.V., Raposo, G., Tian, J.M., Karin, M., Niemann, H., Louvard, D. (1998). A novel tetanus neurotoxin-insensitive vesicle-associated membrane protein in SNARE complexes of the apical plasma membrane of epithelial cells. Mol. Biol. Cell 9, 1437–1448.

Gupton, S.L., and Gertler, F.B. (2010). Integrin signaling switches the cytoskeletal and exocytic machinery that drives neuritogenesis. Dev. Cell 18, 725–736.

Hahn, U. (2012). A Studentized Permutation Test for the Comparison of Spatial Point Patterns. J. Am. Stat. Assoc. 107, 754–764.

Heuser, J.E., and Reese, T.S. (1973). Evidence for recycling of synaptic vesicle membrane during transmitter release at the frog neuromuscular junction. J. Cell Biol. 57, 315–344.

Jaqaman, K., Loerke, D., Mettlen, M., Kuwata, H., Grinstein, S., Schmid, S.L., and Danuser, G. (2008). Robust single-particle tracking in live-cell time-lapse sequences. Nat. Methods 5, 695–702.

Kalman, R.E. (1960). A New Approach to Linear Filtering and Prediction Problems. Int. J. Eng. Trans. A 82, 35.

Link, E., Edelmann, L., Chou, J.H., Binz, T., Yamasaki, S., Eisel, U., Baumert, M., Südhof, T.C., Niemann, H., and Jahn, R. (1992). Tetanus toxin action: inhibition of neurotransmitter release linked to synaptobrevin proteolysis. Biochem. Biophys. Res. Commun. 189, 1017–1023.

Li, Y., Chin, L.-S., Weigel, C., and Li, L. (2001). Spring, a Novel RING Finger Protein That Regulates Synaptic Vesicle Exocytosis. J. Biol. Chem. 276, 40824–40833.

Martinez-Arca, S., Alberts, P., Zahraoui, A., Louvard, D., Galli, T. (2000). Role of tetanus neurotoxin insensitive vesicle-associated membrane protein (TI-VAMP) in vesicular transport mediating neurite outgrowth. J. Cell Biol. 149, 889–900.

Matov, A., Applegate, K., Kumar, P., Thoma, C., Krek, W., Danuser, G., and Wittmann, T. (2010). Analysis of microtubule dynamic instability using a plus-end growth marker. Nat. Methods 7, 761–768.

Mauch, D.H., Nägler, K., Schumacher, S., Göritz, C., Müller, E.C., Otto, A., and Pfrieger, F.W. (2001). CNS synaptogenesis promoted by glia-derived cholesterol. Science 294, 1354–1357.

McMahon, H.T., Ushkaryov, Y.A., Edelmann, L., Link, E., Binz, T., Niemann, H., Jahn, R., Südhof, T.C., (1993). Cellubrevin is a ubiquitous tetanus-toxin substrate homologous to a putative synaptic vesicle fusion protein. Nature 364, 346–349.

Menon, S., Boyer, N.P., Winkle, C.C., McClain, L.M., Hanlin, C.C., Pandey, D., Rothenfußer, S., Taylor, A.M., and Gupton, S.L. (2015). The E3 Ubiquitin Ligase TRIM9 Is a Filopodia Off Switch Required for Netrin-Dependent Axon Guidance. Dev. Cell 35, 698–712.

Miesenböck, G., De Angelis, D.A., and Rothman, J.E. (1998). Visualizing secretion and synaptic transmission with pH-sensitive green fluorescent proteins. Nature 394, 192–195.

Mostov, K.E., Verges, M., and Altschuler, Y. (2000). Membrane traffic in polarized epithelial cells. Curr. Opin. Cell Biol. 12, 483–490.

Oishi, Y., Arakawa, T., Tanimura, A., Itakura, M., Takahashi, M., Tajima, Y., Mizoguchi, I., and Takuma, T. (2006). Role of VAMP-2, VAMP-7, and VAMP-8 in constitutive exocytosis from HSY cells. Histochem. Cell Biol. 125, 273–281.

Okabe, A., and Yamada, I. (2001). The K-Function Method on a Network and its computational implementation. Geogr. Anal. 33, 270–290.

Paul, L.K., Brown, W.S., Adolphs, R., Tyszka, J.M., Richards, L.J., Mukherjee, P., and Sherr, E.H. (2007). Agenesis of the corpus callosum: genetic, developmental and functional aspects of connectivity. Nat. Rev. Neurosci. 8, 287–299.

Pawley, J. (1990). Fundamental Limits in Confocal Microscopy, in: Handbook of Biological Confocal Microscopy. pp. 15–26.

Pearse, B.M. (1976). Clathrin: a unique protein associated with intracellular transfer of membrane by coated vesicles. Proc. Natl. Acad. Sci. U. S. A. 73, 1255–1259.

Petersen, J.D., Kaech, S., and Banker, G. (2014). Selective microtubule-based transport of dendritic membrane proteins arises in concert with axon specification. J. Neurosci. 34, 4135–4147.

Pfenninger, K.H. (2009). Plasma membrane expansion: a neuron’s Herculean task. Nat. Rev. Neurosci. 10, 251–261.

Pfenninger, K.H., and Friedman, L.B. (1993). Sites of plasmalemmal expansion in growth cones. Brain Res. Dev. Brain Res. 71, 181–192.

Plooster, M., Menon, S., Winkle, C.C., Urbina, F.L., Monkiewicz, C., Phend, K.D., Weinberg, R.J., Gupton, S.L. (2017). TRIM9-dependent ubiquitination of DCC constrains kinase signaling, exocytosis, and axon branching. Mol. Biol. Cell 28, 2374–2385.

Popov, S., Brown, A., and Poo, M. (1993). Forward plasma membrane flow in growing nerve processes. Science 259, 244–246.

Protopopov, V., Govindan, B., Novick, P., and Gerst, J.E. (1993). Homologs of the synaptobrevin/VAMP family of synaptic vesicle proteins function on the late secretory pathway in S. cerevisiae. Cell 74, 855–861.

Ripley, B.D. (1976). The Second-Order Analysis of Stationary Point Processes. J. Appl. Probab. 13, 255.

Ros, O., Cotrufo, T., Martínez-Mármol, R., and Soriano, E. (2015). Regulation of patterned dynamics of local exocytosis in growth cones by netrin-1. J. Neurosci. 35, 5156–5170.

Schmoranzer, J., Kreitzer, G., and Simon, S.M. (2003). Migrating fibroblasts perform polarized, microtubule-dependent exocytosis towards the leading edge. J. Cell Sci. 116, 4513–4519.

Sebastian, R., Diaz, M.-E., Ayala, G., Letinic, K., Moncho-Bogani, J., and Toomre, D. (2006). Spatio-temporal analysis of constitutive exocytosis in epithelial cells. IEEE/ACM Trans. Comput. Biol. Bioinform. 3, 17–32.

Serafini, T., Kennedy, T.E., Gaiko, M.J., Mirzayan, C., Jessell, T.M., and Tessier-Lavigne, M. (1994). The netrins define a family of axon outgrowth-promoting proteins homologous to C. elegans UNC-6. Cell 78, 409–424.

Shaner, N.C., Lin, M.Z., McKeown, M.R., Steinbach, P.A., Hazelwood, K.L., Davidson, M.W., and Tsien, R.Y. (2008). Improving the photostability of bright monomeric orange and red fluorescent proteins. Nat. Methods 5, 545–551.

Söllner, T., Bennett, M.K., Whiteheart, S.W., Scheller, R.H., and Rothman, J.E. (1993). A protein assembly-disassembly pathway in vitro that may correspond to sequential steps of synaptic vesicle docking, activation, and fusion. Cell 75, 409–418.

Stefan, C.J., Manford, A.G., and Emr, S.D. (2013). ER-PM connections: sites of information transfer and inter-organelle communication. Curr. Opin. Cell Biol. 25, 434–442.

Taylor, A.M., Menon, S., and Gupton, S.L. (2015). Passive microfluidic chamber for long-term imaging of axon guidance in response to soluble gradients. Lab Chip 15, 2781–2789.

Tojima, T., Akiyama, H., Itofusa, R., Li, Y., Katayama, H., Miyawaki, A., and Kamiguchi, H. (2007). Attractive axon guidance involves asymmetric membrane transport and exocytosis in the growth cone. Nat. Neurosci. 10, 58–66.

Tojima, T., Itofusa, R., and Kamiguchi, H. (2014). Steering neuronal growth cones by shifting the imbalance between exocytosis and endocytosis. J. Neurosci. 34, 7165–7178.

Viesselmann, C., Ballweg, J., Lumbard, D., Dent, E.W., 2011. Nucleofection and primary culture of embryonic mouse hippocampal and cortical neurons. J. Vis. Exp. doi:10.3791/2373

Vincent, L., (1993). Morphological grayscale reconstruction in image analysis: applications and efficient algorithms. IEEE Trans Image Process, 2(2):176–201.

Waters, J.C., (2009). Accuracy and precision in quantitative fluorescence microscopy. J. Exp. Med. 206, i15–i15.

Winckler, B., Forscher, P., and Mellman, I., (1999). A diffusion barrier maintains distribution of membrane proteins in polarized neurons. Nature 397, 698–701.

Winkle, C.C., McClain, L.M., Valtschanoff, J.G., Park, C.S., Maglione, C., and Gupton, S.L. (2014). A novel Netrin-1–sensitive mechanism promotes local SNARE-mediated exocytosis during axon branching. J. Cell Biol. 205, 217–232.

Winkle, C.C., Olsen, R.H.J., Kim, H., Moy, S.S., Song, J., and Gupton, S.L. (2016). Trim9 Deletion Alters the Morphogenesis of Developing and Adult-Born Hippocampal Neurons and Impairs Spatial Learning and Memory. J. Neurosci. 36, 4940–4958.

Yuan, T., J. Lu, J. Zhang, Y. Zhang, and L. Chen. (2015). Spatiotemporal Detection and Analysis of Exocytosis Reveal Fusion “Hotspots” Organized by the Cytoskeleton in Endocrine Cells. Biophys J. 108:251–260.

